# Regulation of PDGF-BB Signaling in Placental Pericytes by Soluble PDGFRβ Isoforms: Implications for Fetoplacental Vascular Development

**DOI:** 10.64898/2026.03.24.713995

**Authors:** Audra Barnes, Emily C. Duggan, Reese Dunkenberger, Chloe Lessard, Christopher Cosma, Chase Steele, Sarah V. Taylor, Megan D. Whitham, Allison R. Durica, John C. Chappell

## Abstract

Vascular remodeling within the developing fetus and placenta is essential for supporting the growth and function of emerging tissues and organs. Pericytes (PCs) play a central role in stabilizing and maturing microvascular networks by extending along endothelial cells (ECs) and reinforcing vessel integrity. In the placenta, as in other organs, PC–EC communication is mediated in part by platelet-derived growth factor-BB (PDGF-BB) signaling, which governs PC differentiation, proliferation, migration, and survival, ultimately enabling their recruitment and retention along capillaries. In this study, we identified progressive PC investment along feto-placental capillaries in both murine and human tissues across gestation, supported by morphological and molecular evidence. Placental PCs displayed phenotypic heterogeneity comparable to that observed in the brain and heart, suggesting conserved diversity across organ systems. In addition to characterizing PC dynamics, we examined the expression of recently identified soluble PDGF Receptor-β (sPDGFRβ) isoforms. These variants were detected at the protein and transcript levels in mouse and human placentas, as well as in a murine trophoblast-embryonic stem cell (TESC) differentiation model that recapitulates aspects of early placental vascular development. Within this model, sPDGFRβ expression was independent of ADAM10 activity and exogenous growth factors during early vessel formation but was markedly upregulated during hypoxia. To assess how elevated sPDGFRβ might influence PDGF-BB signaling, we exposed TESCl-derived vascular networks to excess PDGF-BB with or without a sPDGFRβ mimetic. PDGF-BB alone reduced full-length PDGFRβ levels while increasing receptor phosphorylation, consistent with known ligand-induced regulatory mechanisms. Inclusion of the sPDGFRβ mimetic shifted these responses toward baseline, suggesting a potential modulatory or feedback role for soluble receptor variants. Together, these findings demonstrate that PCs are progressively recruited to placental capillaries and exhibit diverse phenotypes during development, and that soluble PDGFRβ isoforms may modulate PDGF-BB signaling in a manner sensitive to oxygen tension. Understanding these mechanisms provides insight into the regulation of placental vascular maturation and may inform strategies to improve human health by targeting disorders rooted in impaired placental development.

## INTRODUCTION

Placental development is fundamental to embryogenesis and exerts lasting effects on postnatal health for both mom and baby [1, 2]. A functional placental vascular network must form, expand, and mature appropriately to ensure adequate nutrient and oxygen delivery to the fetus [3]. In cases of placental insufficiency, often characterized by impaired perfusion and oxygenation, fetoplacental vessels often remain in a pathological state of remodeling, disrupting the balance of molecular signals that regulate vascular growth and stability [4, 5]. This is seen in preeclampsia, where excess release of soluble vascular endothelial growth factor receptor-1 (sVEGFR1, or sFlt-1 in mice) into maternal circulation contributes to systemic hypertension and heightened perinatal risk [6–8]. Elucidating molecular drivers of placental vascular dysfunction remains a critical step toward mitigating both immediate and long-term health consequences of placental insufficiency and disease. Placental insufficiency for example is a major cause of fetal growth restriction and preterm birth, often rooted in disrupted vascular development [4, 5]. Normally, fetoplacental vessels expand and remodel to support increasing metabolic needs, but during insufficiency, this process stalls or maladapts [9, 10]. Hypoxia and oxidative stress in the placenta alter signaling pathways that control vascular remodeling, resulting in leaky, immature, or poorly perfused vessels [9, 11–13]. This dysfunction impairs nutrient and gas exchange, threatening fetal development and survival [14, 15]. Thus, identifying the mechanistic determinants of abnormal remodeling is essential for developing biomarkers and therapeutic interventions to manage and improve placental function in at-risk pregnancies.

During fetal development, pericytes (PCs) stabilize and mature the microvasculature of the placenta by wrapping around endothelial cells (ECs) and contributing to vessel morphology and integrity [16–19]. In human placental terminal villi, PC “foot processes” preferentially cover endothelial junctions distant from trophoblasts (TBs), which likely affects fluid and molecule diffusion across the capillary wall [20, 21]. PC subsets also express contractile and signaling molecules that integrate with EC-derived cues to regulate vessel diameter, barrier properties, and vascular remodeling [22–25]. Studies using in vitro and 3D coculture models have implicated PC–EC dysfunction in reduced vessel perfusion and growth under pathological conditions [17]. While PCs are gaining attention for their importance in placental development and function, there is more to uncover in identifying their roles in normal fetoplacental vascular formation and in protecting against vascular pathology in complex pregnancies.

Pericyte recruitment and retention within the capillary wall requires the platelet-derived growth factor-BB (PDGF-BB) pathway, as this signaling axis is essential for orchestrating PC differentiation, proliferation, migration, and survival [18, 26–35]. Deficient PDGF-BB signaling leads to PC loss from the capillary wall, causing vessel breach, hemorrhage, and general dysmorphogenesis [28, 30], notably in the placenta [18, 36]. In contrast, excess PDGF Receptor-β (PDGFRβ) signals induce PC hyper-proliferation and myofibroblast transition, leading to vascular anomalies and fibrotic accumulation, evident in patients suffering from *PDGFRB* gain-of-function mutations [37–42] and in placentas of murine activating mutation models [43]. These observations suggest that PDGF-BB signaling must be maintained in a specific range during vessel formation and maturation to promote sufficient PC investment into nascent capillaries to support their function in the placenta and elsewhere in the human body. PDGF-BB signaling has been studied for some time, yet as shown by recent identification of soluble PDGFRβ (sPDGFRβ) isoforms by our lab and others [44–47], we are still learning how this pathway coordinates cellular functions across many tissues in health and disease.

Soluble receptor isoforms have been identified across numerous signaling pathways [48]. In the epidermal growth factor (EGF) pathway, for instance, certain receptors undergo ectodomain shedding, releasing soluble fragments that can modulate signaling by the full-length receptor [49]. This phenomenon is not unique to EGF, as several receptor tyrosine kinase (RTK) pathways—including fibroblast growth factor (FGF) and VEGF-A signaling networks—involve soluble receptors through alternative mRNA splicing [50–55], a process influenced in part by RNA-binding proteins [56]. A well-studied example is sVEGFR1/sFlt-1, a truncated splice variant that acts as a decoy receptor or ligand sink to regulate VEGF-A bioavailability and, in turn, signaling capacity [57–59]. As mentioned above, the PDGF-BB system also appears to incorporate soluble isoforms of the full-length PDGFRβ. These may arise from truncated transcripts and possibly enzymatic cleavage via a disintegrin and metalloproteinase domain-containing protein 10 (ADAM10), though proteolysis may be context-dependent [44, 46, 47]. Cleaved sPDGFRβ has been proposed as a degradation product associated with PC dysfunction or loss [47], rather than a molecule with a defined mechanistic role. We recently found that alternatively spliced mRNA transcripts are generated by murine and human cells under standard culture conditions, with murine brain lysates also containing truncated sPDGFRβ isoforms on both the protein and mRNA levels during normal development [45]. Thus, greater insight will be needed to shed light on the regulatory mechanisms underlying sPDGFRβ production in health and disease, and what specific roles sPDGFRβ isoforms might play in modulating PDGF-BB signaling in relevant cell types including capillary PCs.

To better understand placental PC biology and mechanistic determinants within the PDGF-BB pathway, we identified both morphological and molecular evidence from murine and human placental tissues, demonstrating a progressive increase in PC coverage of feto-placental capillaries across gestation. Placental PCs also displayed phenotypic diversity mirroring organs such as the brain, heart, and lungs [60–62]. In addition to characterizing mechanisms that regulate vessel formation and PC expansion, we detected recently described soluble isoforms of PDGFRβ at the protein and transcript levels in both mouse and human placentas. These soluble variants were also expressed in a murine stem-cell–based differentiation model that recapitulates key aspects of early placental vascular development, including vessel formation, PC involvement, and TB lineage emergence. Using this TB and embryonic stem cell (TESC) model, we found that sPDGFRβ production during early vascular development did not appear to depend on ADAM10 activity or exogenous growth factor exposure. Hypoxia, however, markedly increased the abundance of sPDGFRβ isoforms, consistent with previous studies [45]. To explore how elevated sPDGFRβ might influence PDGF-BB signaling, we treated TESC-derived vessels with excess PDGF-BB in the presence or absence of an sPDGFRβ mimetic. PDGF-BB alone reduced full-length PDGFRβ and increased receptor phosphorylation, consistent with established regulatory mechanisms [63, 64], whereas co-treatment with the sPDGFRβ mimetic shifted these responses toward baseline, aligning with previous studies [33, 65, 66]. Collectively, our findings indicate that PCs are progressively recruited to capillaries at the maternal-fetal interface during murine and human placental development and exhibit heterogeneity comparable to that seen in other organs. Moreover, our data suggest that soluble PDGFRβ isoforms may modulate full-length PDGFRβ signaling in this environment, potentially independent of ADAM10 activity or growth factor availability but responsive to reduced oxygen tension.

## METHODS AND MATERIALS

### In Vivo Animal and Human Studies

In the current study, we utilized placenta tissue samples from C57BL/6 and transgenic mice harboring a gene construct labeling capillary pericytes expressing Ng2/Cspg4 (e.g. oligodendrocyte precursors), specifically with the DsRed2 fluorophore (STOCK Tg(Cspg4-DsRed.T1)1Akik/J, NG2DsRedBAC, The Jackson Laboratory, Strain #008241) [67]. In total, 15 adult female mice and 40 pups were used for this study between the ages of 3- and 6-months old, under approval granted by the Institutional Animal Care and Use Committee (IACUC), via protocols #23-167 and #25-081. All animals were purchased from The Jackson Laboratory or generated within our in-house animal colony. All experiments involving animal use were performed following review and approval from the IACUC at Virginia Tech. All experimental protocols were reviewed and approved by Virginia Tech Veterinary Staff and the IACUC. The Virginia Tech NIH/PHS Animal Welfare Assurance Number is A-3208-01. All methods were performed in accordance with the relevant guidelines and regulations provided by these entities. This study was conducted, and is reported, in accordance with ARRIVE guidelines. We also utilized placenta tissue samples from human patients who provided their full consent for the specific uses in the study under the IRB Protocols #25-1971 and #21-1345, which were reviewed and approved by the Carilion Clinic Human Research Protections Office (HRPO) and Institutional Review Board (IRB). Samples included for the current study were from uncomplicated, full-term pregnancies (>37 weeks of gestation) of mothers 18 years of age or older; exclusion criteria included diagnosis of gestational hypertension, preeclampsia, gestational diabetes, placental abruption, history of preterm delivery resulting in long-term complications for the child, chorioamnionitis at delivery.

### In Vitro Cell Culture Studies

Trophoblast and embryonic stem cells (TESCs) collected previously from WT or “double-reporter” (DR – *Flk1^eGFP/+^; Cspg4^DsRed/+^*) [68] mice were maintained as undifferentiated cell lines via exposure to leukemia inhibitory factor (LIF). Upon LIF removal (i.e. differentiation day 0, dd0), TESCs were allowed to aggregate into spheroids that were transferred into a 10 cm^2^ plate or individual wells of a 24-well plate for single-spheroid culture for 7 to 12 days of differentiation, depending on the experimental designation. For bulk RNA sequencing of dd0, dd7, or dd10 cells following fluorescence-activated cell sorting (FACS), we collected and analyzed cells expressing eGFP only, DsRed only, or cells lacking fluorescence per the detailed protocol provided in Payne et al. [68]. For western blot analysis of sPDGFRβ protein levels, cells were washed with PBS and lysed to collect protein at dd8, dd10, and dd12. For ADAM10 inhibitor experiments, TESCs were treated at dd8 and dd10 during media changes with GI-254023X (10uM) or DMSO (equivalent volume, vehicle control), with protein collected at dd12 for western blot analysis. For hypoxia experiments, TESCs were differentiated for 10-days under standard culture conditions (ambient O_2_ levels + 5% CO_2_). At dd10, cells were moved to 3% O_2_ (5% CO_2_ + 92% N_2_) for 12- or 48-hrs, at which time protein was collected for western blot analysis. For growth factor treatment experiments, we differentiated TESCs under standard media conditions until dd8, replacing this media with one of the following: (1) endothelial growth media (EGM), basal form (Lonza) plus 10% fetal bovine serum (FBS), (2) EGM + 10% FBS + VEGF-A (10ng/mL, PeproTech), (3) EGM + 10% FBS + PDGF-BB (10ng/mL, PeproTech), or (4) EGM + 10% FBS + EGM Kit Supplements (Lonza) + PDGF-BB (10ng/mL). For soluble PDGFRβ mimetic experiments, we cultured TESCs under standard conditions for spontaneous differentiation until day 8, at which time we switch media to one of the following: (1) baseline media with EGM basal form media + 10% FBS, (2) EGM + 10% FBS + PDGF-BB (10ng/mL), and (3) EGM + 10% FBS + PDGF-BB (10ng/mL) + soluble PDGFRβ-Fc chimera (sPDGFRβ-Fc or sRβ-Fc, R&D Systems).

### Placenta Slice and Differentiated TESC Immunolabeling, Confocal Imaging, and Image Quantification

Human (full-term) and murine (embryonic day 11.5, E14.5, E18.5) placenta slices used for immunostaining and confocal imaging were fixed in 4% paraformaldehyde (PFA) at 4°C – 4 days for human tissues, overnight for murine – and then washed and stored in PBS. Differentiated TESCs were fixed with 4% PFA for 15-mins before PBS wash and transfer to a glass-bottom plate. Non-specific antigens within each cryostat-generated section or culture were blocked using PBS-T (0.5% Triton-X 100 in PBS) with 5% bovine serum albumin (BSA) for 1-hr. Samples were then incubated in PBS-T containing primary antibodies (all at 1:200) overnight at 4°C: PECAM-1 (R&D Systems), NG2 (Millipore), PDGFRβ (ThermoFisher), αSMA (Sigma-Millipore), or TROMA-I (Developmental Studies Hybridoma Bank). They were washed in PBS-T (2x, 10-min each wash, at room temperature) followed by 1x PBS (1x, 5-min, at room temperature) and incubated with appropriate secondary antibodies (all at 1:400) for 3-hrs at room temperature, as needed. Nuclei were stained with DAPI (1:1000) for 10-mins at room temperature. Slices were mounted on glass slides with glass coverslips and imaged using a Zeiss LSM 880 confocal microscope with volumetric, z-stack images acquired. Confocal images were analyzed for the signals indicated using QuPath® [69]. Raw image analysis files are available upon request.

### Quantitative RT-PCR

Messenger RNA transcripts were extracted from E9.5, E14.5, and E18.5 murine placentas in Trizol^TM^, and dd8, dd10, dd12 TESCs using RNA Lysis Buffer (Zymo®), followed by precipitation via Zymo® column-based kits with on-column DNase treatment using the reagent from the Zymo® kit and TURBO DNase™ (removing genomic DNA) and resuspension in RNase-free water. mRNA purity and concentration were assessed via a Nanodrop^TM^ system. cDNA synthesis was performed using a reverse transcription kit (High-Capacity cDNA Reverse Transcription Kit, Applied Biosystems), and qRT-PCR was conducted using standard, “best coverage” TaqMan^TM^ assays for the probes indicated on an Applied Biosystems QuantStudio6-Flex® platform. Custom probes were designed and validated to quantify the soluble *Pdgfrβ* intron4 (i4) and intron10 (i10) splice variants: *intron4* with Forward Primer—TACGTCTACAGCCTCCAG, Reverse Primer—TAGTGAACTGATGGGCTATTC, Probe Sequence—CCTCGCCTTCTGGCCCTTCATAGA, and *intron10* with Forward Primer—AACTCCATGGGTGGAGATTC, Reverse Primer—GTGAGAGTCATCAGAGCCATC, Probe Sequence—TCACCGTGGTCCCACATTGTGAG.

### Western Blot

Protein lysates from human (full-term) and murine (E9.5, E14.5, E18.5) placentas were prepared by resuspending isolated microvessels in RadioImmunoPrecipitation Assay (RIPA) buffer with protease and phosphatase inhibitors, followed by homogenization and centrifugation (15,000×g, 20-min, 4°C). Protein lysates from differentiated TESCs were also collected in RIPA buffer with protease and phosphatase inhibitors and were processed as follows. Protein quantification was performed using the Bradford assay, and samples were resolved via SDS-PAGE on 4–12% Bis-Tris gels. Proteins were transferred to polyvinylidene difluoride (PVDF) membranes and probed with primary antibodies overnight at 4°C for PDGFRβ (1:500, all isoforms, R&D Systems), phosphorylated PDGFRβ (1:100, phospho-PDGFRβ, Y751, R&D Systems) and α-tubulin (1:5000, Sigma-Millipore), followed by incubation with secondary antibodies (all double the concentration of primary antibody) for 2-hrs at room temperature. Bands were detected using a BioRad ChemiDoc® system and ImageLab® software, with band density assessed using ImageJ/FIJI application.

## RESULTS

### Pericytes Invest Along Placental Capillaries during Murine and Human Gestation

Pericytes support capillary beds across a broad range of organs and tissues including the brain, heart, and lungs, among many others [60–62]. Within these microvascular networks, PCs appear to exhibit a phenotypic heterogeneity marked by unique morphologies and transcriptional profiles [60, 70, 71]. Recent studies have shown that the feto-placental vasculature also contains PC subtypes [2, 14]. Here, we sought to better understand PC investment dynamics within the context of the developing murine and human placenta. To begin addressing this question, we acquired sections from full-term human placentas from uncomplicated pregnancies, focusing on the chorionic villi containing fetal microvessels that facilitate exchange with maternal blood [72]. We immunolabeled these capillaries for ECs via platelet-endothelial cell adhesion molecule-1 (PECAM-1)/CD31 staining and for mural cells via neural glial antigen-2 (NG2)/*Cspg4* (Figure 1Ai-iii) or α-smooth muscle actin (αSMA)/*Acta2* (Figure 1Aiv-vi) staining. We observed abundant NG2 signal along the abluminal side of numerous vessels, both capillary-sized and larger, while αSMA signal appeared predominantly located on larger vessels with occasional signal along microvessels. A similar staining pattern was observed in immunostained murine E18.5 placenta sections, wherein we found NG2-positive cells along PECAM-1/CD31-labeled microvasculature with a portion containing αSMA-signal (Figure 1Bi-iv, yellow arrows) while others lacked notable staining for this contractile machinery protein (Figure 1Bi-iv, white arrowhead). Immunostaining for the well accepted mural cell marker PDGFRβ revealed cells morphologically consistent with PCs extended along capillary walls (Figure 1Bv-vi). These observations support the idea that PCs are present along microvessels comprising the maternal-fetal interface during placental growth and expansion, exhibiting heterogeneous αSMA expression.

**Figure 1.**
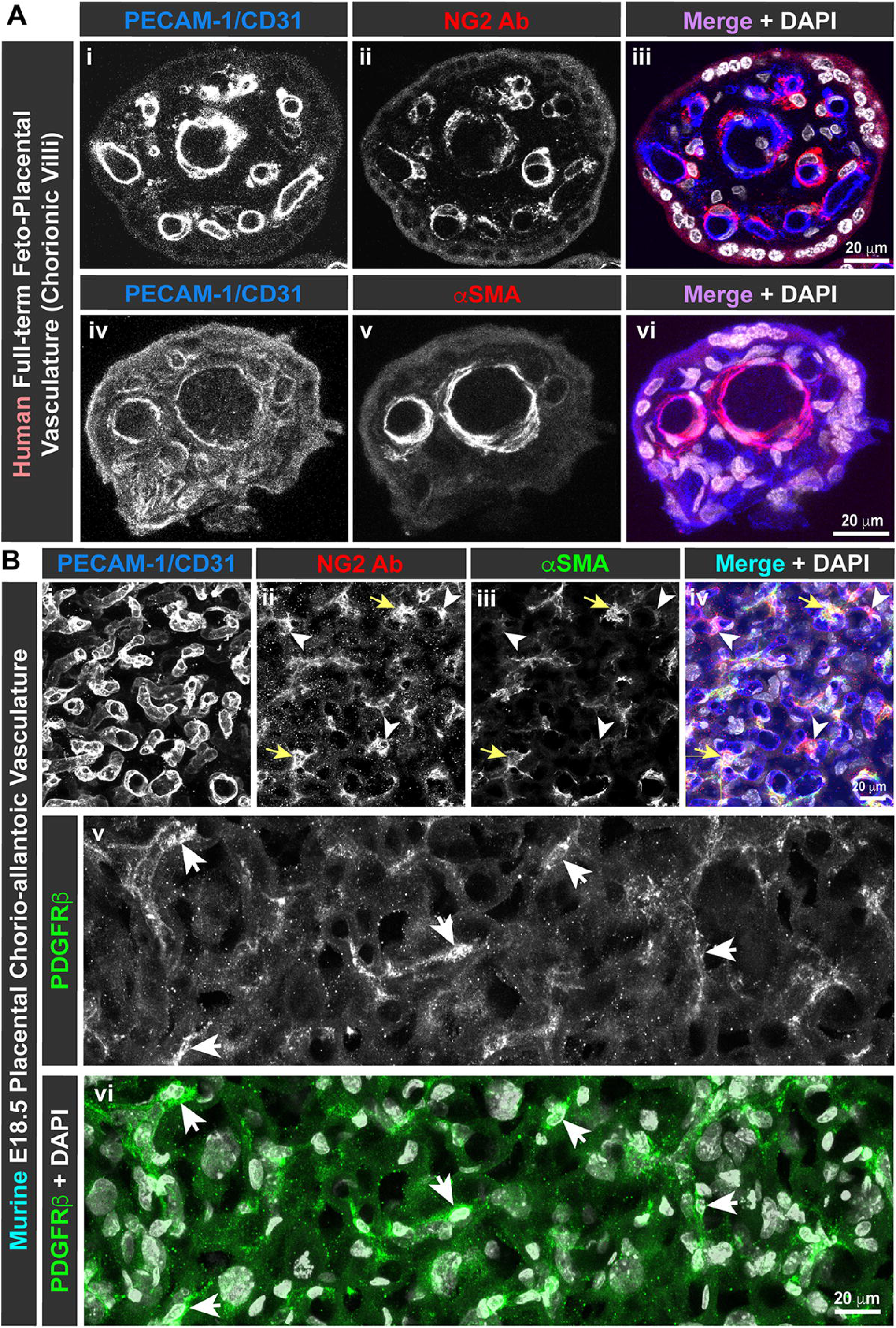
Pericytes Incorporate within Placenta Capillaries of Full-term Human Chorionic Villi and Murine Chorio-allantoic Vasculature. (A) Representative confocal images of human full-term placental sections (chorionic villi) demonstrating immunofluorescent labeling of fetal capillary endothelial cells via PECAM-1/CD31 antibody (i & iv; blue in iii & vi) and perivascular mural cells by NG2 (ii; red in iii) or αSMA (v; red in vi) antibodies. Merged overlays include cell nuclei labeling by DAPI (white in iii & vi). Note extensive pericyte investment within placental vasculature in ii, while αSMA expression appears to be less abundant and spatially heterogeneous in v. Scale bars are 20 mm in iii and vi. (B) Representative confocal images of murine E18.5 placental sections (chorio-allantoic vasculature within the labyrinth zone) demonstrating immunofluorescent labeling of fetal capillary endothelial cells via PECAM-1/CD31 antibody (i; blue in vi) and perivascular mural cells by NG2 antibody (ii; red in iv) and αSMA antibody (iii; green in iv). Merged overlay includes cell nuclei labeling by DAPI (white in iv). Note extensive pericyte investment within placental vasculature, but αSMA expression appears to be heterogeneous. Yellow arrows denote cells positive for both labels, while white arrowheads indicate NG2-positive cells lacking αSMA antibody signal (ii-iv). PDGFRβ immunostaining (v; green in vi) is consistent with pericyte morphology (white arrows) and soluble isoforms distributed more broadly. Merged overlay (vi) includes cell nuclei labeling by DAPI (white). Scale bars are 20 mm in iv and vi.

To better characterize the time-course for PC investment along the placental capillary wall, we collected murine placentas at early (embryonic day 9.5 (E9.5)/E11.5), mid- (E14.5), and late (E18.5) gestation to capture chorio-allantoic vascular remodeling dynamics during each phase [1–3]. Placenta sections from each time point were immunostained for PECAM-1/CD31 to visualize the emerging endothelium, with PCs and their progenitors identified by NG2 immunolabeling (E11.5 & E14.5, Figure 2Ai-ix) or DsRed reporter signal via the *NG2/Cspg4^DsRed/+^* construct [67] (E18.5, NG2:DsRed, Figure 2Ax-xii). During early murine gestation, NG2-positive cells were observed in the context of cells differentiating within the hemangioblast lineage, specifically located along primitive and maturing vascular structures labeled for PECAM-1/CD31 (Figure 2Ai-vi). By E14.5, PCs appeared along and around a vast majority of murine placental capillaries (Figure 2Avii-ix), a trend that was also found at E18.5 (Figure 2Ax-xii). We also noted a subset of NG2-positive PCs within E18.5 placenta vessel networks extending processes across several vessels (Figure 2Ax-xii), consistent with a “cross-bridging” phenotype previously described in other adult murine tissues [61, 73]. PC density increased dramatically over these early placentogenesis stages (Figure 2B), though it appeared to match EC expansion and capillary formation (Supplemental Figure 1). To compare these morphological observations with transcriptional changes, we collected murine placenta mRNA at early (E9.5), mid- (E14.5), and late (E18.5) gestation and analyzed relative transcript levels of PC-related genes by qRT-PCR. *Cspg4* (encoding NG2 protein) and *Acta2* (encoding αSMA) transcripts increased significantly between E9.5 and E14.5 with both plateauing from E14.5 to E18.5, though remaining elevated relative to E9.5 (Figure 2C-D). Full-length *Pdgfr*β expression rose from E9.5 to E14.5, displaying a significant increase from E14.5 to E18.5 (Figure 2E). The mural cell-derived ECM component vitronectin (*Vtn*) [25]experienced a significant transcriptional increase from E9.5 to E14.5 but then decreased significantly by E18.5, though not returning to E9.5 levels (Figure 2F). *Notch3* expression [74–76] followed a similar trend as *Vtn* across the timepoints collected, though statistically significant differences were not found (Figure 2G). Overall, these morphological and transcriptional observations further establish the importance of PC contributions to the placental chorio-allantoic vasculature throughout murine and human gestation.

**Figure 2.**
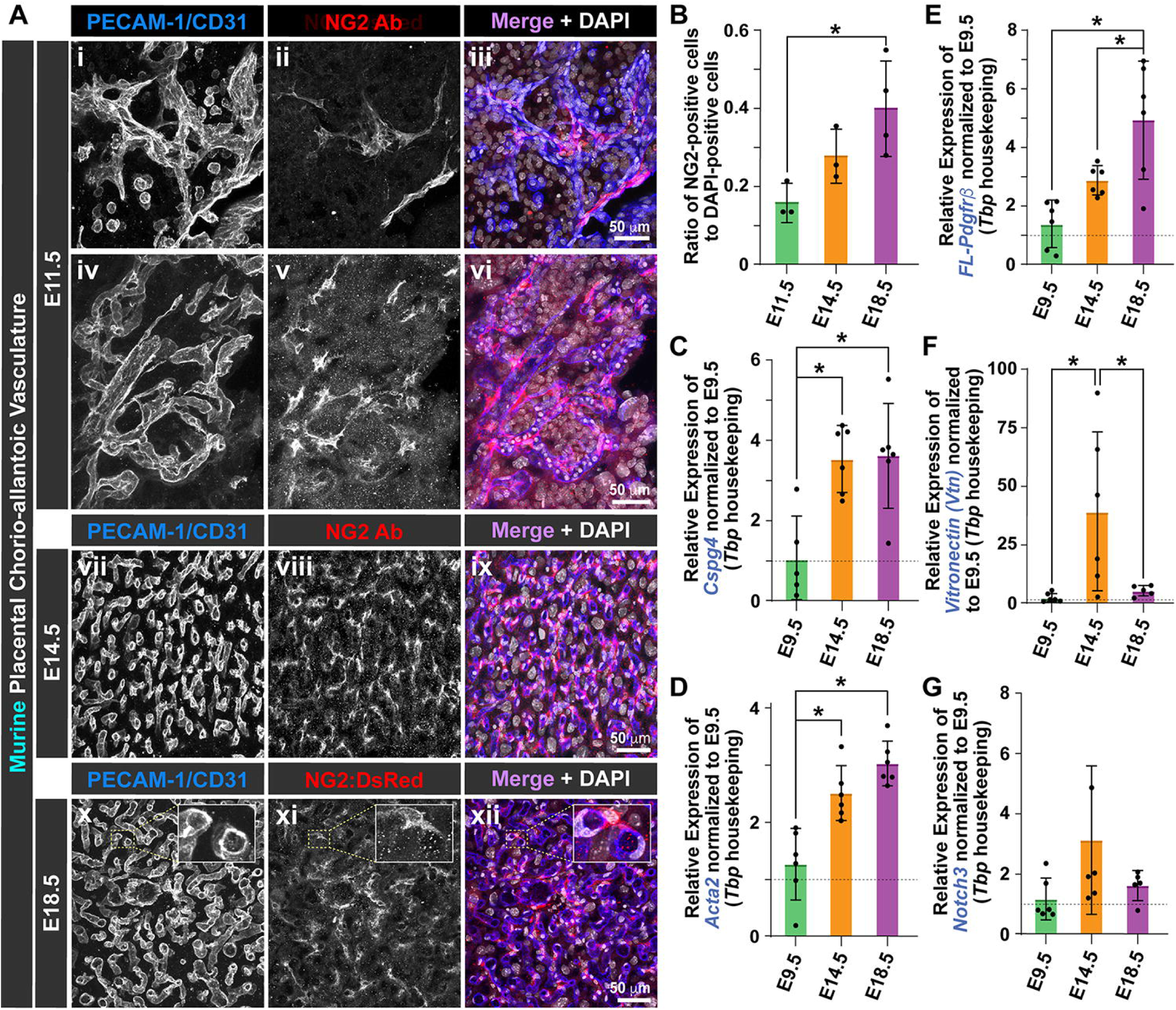
Pericytes Progressively Invest within the Murine Chorio-allantoic Vasculature, Mirroring Perivascular Cell Gene Expression during Placenta Formation and Expansion. (A) Representative confocal images of murine placental sections (chorio-allantoic vessels within the developing labyrinth zone) demonstrating reporter and immunofluorescent labeling of fetal capillary endothelial cells via PECAM-1/CD31 antibody at E11.5 (i & iv; blue in iii & vi), E14.5 (vii; blue in ix), and E18.5 (x; blue in xii) and perivascular mural cells at corresponding timepoints by NG2 antibody (ii, v, & viii; red in iii, vi, & ix) or *NG2/Cspg4^DsRed/+^* reporter (xi; red in xii). Merged overlays include cell nuclei labeling by DAPI (white in iii, vi, ix, & xii). Scale bars are 50 mm in iii, vi, ix, and xii. (B) Graph of NG2-positive cells normalized to all DAPI-positive cells for a given image from the indicated time points, E11.5 (n=3, green bar), E14.5 (n=3, orange bar), and E18.5 (n=4, purple bar). Individual data points shown, and errors bars are standard deviation. *p≤0.05 by one-way ANOVA with post-hoc Tukey test for comparison indicated. *Note in (C-G), for each gene expression graph shown: all data were normalized to E9.5 as indicated by the dotted line, Tbp served as the housekeeping gene, individual data points were plotted for each embryonic day with averages represented by bars: E9.5--green, E14.5--orange, E18.5--purple, error bars represent standard error of the mean, and *p*≤*0.05 by one-way ANOVA with post-hoc Tukey test for comparisons indicated.* (C) Graph of relative *Cspg4* expression within the murine placenta at E9.5 (n=5), E14.5 (n=6), and E18.5 (n=6). (D) Graph of relative *Acta2 (αSMA)* expression within murine placenta at E9.5 (n=6), E14.5 (n=6), and E18.5 (n=6). (E) Graph of relative *Full Length (FL)-Pdgfrb* expression within murine placenta at E9.5 (n=6), E14.5 (n=6), and E18.5 (n=6). (F) Graph of relative *Vitronectin (Vtn)* expression within murine placenta at E9.5 (n=6), E14.5 (n=6), and E18.5 (n=5). (G) Graph of relative *Notch3* expression within murine placenta at E9.5 (n=6), E14.5 (n=5), and E18.5 (n=5).

### Soluble PDGF Receptor-β Isoforms Emerge during Vessel Formation in Murine and Human Placentas and in a Murine Stem Cell Model of Placental Vascular Development

Placental vascular development requires an intricate and complex balance of pro- and anti-angiogenic molecular cues that align with specific phases of placenta growth and maturation [10, 77]. Here, we focused on the VEGF-A and PDGF-BB pathways to better understand how EC and PC signaling and negative feedback regulators might be coordinated during this process. Specifically, we applied qRT-PCR to mRNA collected from E9.5, E14.5, and E18.5 murine placentas to assess gene expression of molecular components within these pathways. Transcripts for *Vegfa* increased significantly from E9.5 to E14.5 and remained unchanged from E14.5 to E18.5 (Figure 3A). The VEGF-A receptor Flt-1, which functions largely as a “decoy” receptor to modulate VEGF-A ligand bioavailability [57–59], increased significantly on the transcriptional level from E9.5 to E14.5, then decreased from E14.5 to E18.5 (Figure 3B). The primary VEGF-A signaling receptor in ECs *Flk-1* (or *Kdr*) displayed an increase from E9.5 to E14.5 and then a decrease from E14.5 to E18.5, but neither change was statistically significant (Figure 3C). In contrast, Placental Growth Factor (PlGF) transcripts decreased from E9.5 to E14.5 and again from E14.5 to E18.5, reaching a significantly lower level of expression at E18.5 compared to E9.5 (Figure 3D). Expression of PDGF-B (*Pdgfb*) rose from E9.5 to E14.5, with a significant increase from E14.5 to E18.5 (Figure 3E). A similar trend was observed for transcripts encoding the intron 4 truncation variant of sPDGFRβ (*sPdgfr*β, i4, Figure 3F).

**Figure 3.**
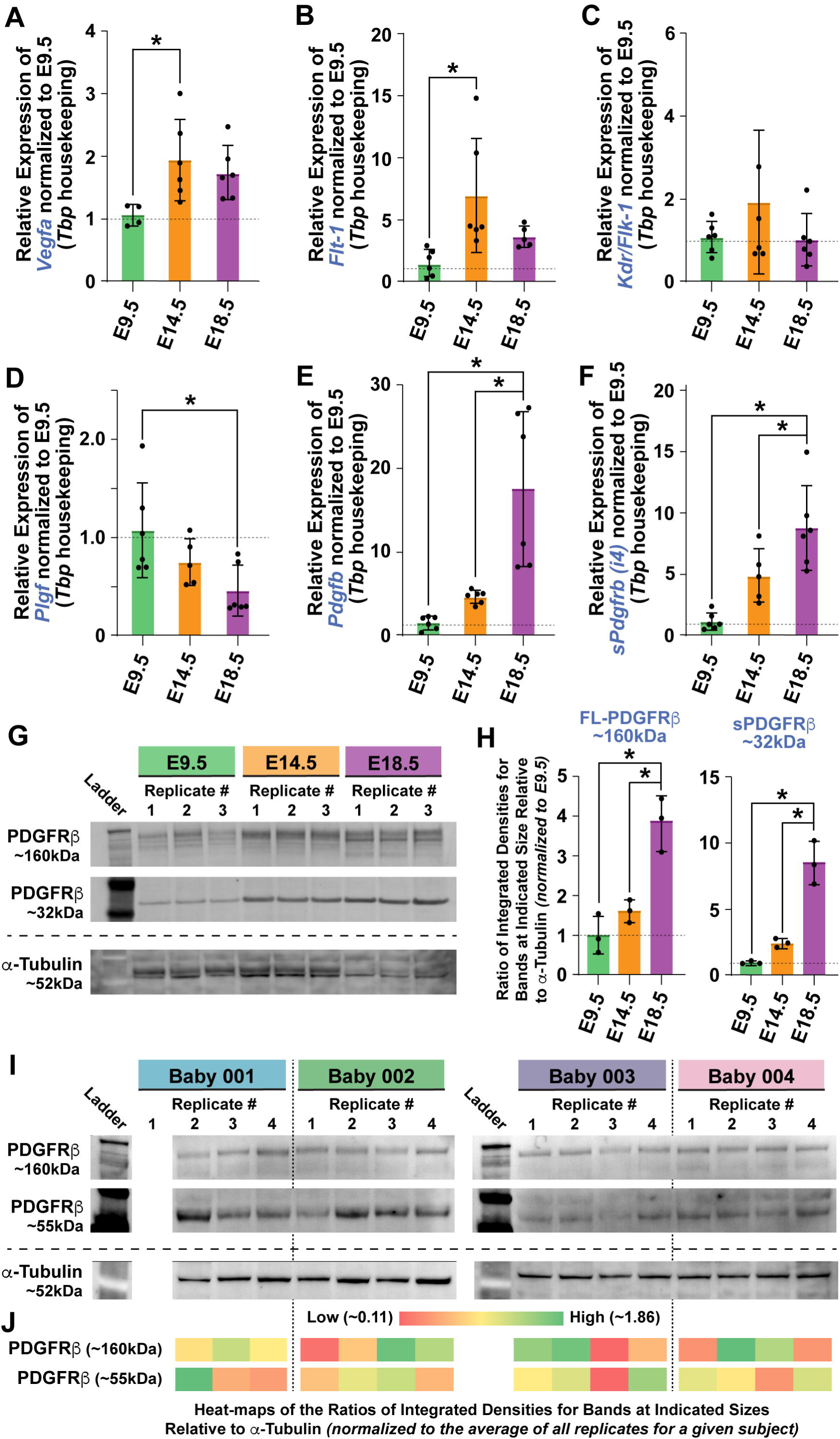
Vascular Remodeling Cues Exhibit Coordinated Transcriptional Changes within the Murine Placenta over Gestation including Soluble PDGF Receptor-β, which Increases on the Protein Level later in Pregnancy and Exhibits Regional Heterogeneity in the Human Placenta. *Note in (A-F), for each gene expression graph shown: all data were normalized to E9.5 as indicated by the dotted line, Tbp served as the housekeeping gene, individual data points were plotted for each embryonic day with averages represented by bars: E9.5--green, E14.5--orange, E18.5--purple, error bars represent standard error of the mean, and *p*≤*0.05 by one-way ANOVA with post-hoc Tukey test for comparisons indicated.* (A) Graph of relative *Vegfa* expression within the murine placenta at E9.5 (n=4), E14.5 (n=6), and E18.5 (n=6). (B) Graph of relative *Flt-1* expression within murine placenta at E9.5 (n=6), E14.5 (n=6), and E18.5 (n=5). (C) Graph of relative *Kdr/Flk-1* expression within murine placenta at E9.5 (n=6), E14.5 (n=5), and E18.5 (n=6). (D) Graph of relative *PlGF* expression within murine placenta at E9.5 (n=6), E14.5 (n=5), and E18.5 (n=6). (E) Graph of relative *Pdgfb* expression within murine placenta at E9.5 (n=6), E14.5 (n=6), and E18.5 (n=6). (F) Graph of relative *soluble Pdgfrb* expression (intron4 truncation variant) within murine placenta at E9.5 (n=6), E14.5 (n=5), and E18.5 (n=6). (G) Representative western blot images of murine placenta protein labeled for PDGFRβ (∼160kDa and ∼32kDa) with a-tubulin (∼52kDa) serving as loading control. Ladder in Lane 1, 3 replicates of E9.5 in Lanes 2-4, 3 replicates of E14.5 in Lanes 5-7, and 3 replicates of E18.5 in Lanes 8-10. *Images were cropped and inverted from original blots, which are provided in Supplemental* Figure 2. (H) Graph of ratios of integrated densities for bands at the indicated sizes relative to α-tubulin (loading control run on same blot). Individual data points are shown with averages represented as bars (E9.5--green, E14.5--orange, E18.5--purple), with dotted gray lines indicating reference to E9.5 samples. *p≤0.05 by one-way ANOVA with post-hoc Tukey test for comparisons indicated. (I) Representative western blot images of human placenta protein labeled for PDGFRβ (∼160kDa and ∼55kDa) with α-tubulin (∼52kDa) serving as loading control. Ladder in Lane 1 of Blots 1 & 2; 3 distinct replicates from Baby 001 in Lanes 3-5, Blot 1 (*Note, Replicate #1 in Lane 2 not shown due to technical error)*; 4 distinct replicates from Baby 002 in Lanes 6-10, Blot 1; 4 distinct replicates from Baby 003 in Lanes 2-5, Blot 2; 4 distinct replicates from Baby 004 in Lanes 6-10, Blot 2. *Images were cropped and inverted from original blots, which are provided in Supplemental* Figures 3-4. (J) Heatmap of the ratios of integrated densities for bands at the indicated sizes relative to α-tubulin (loading control run on same blot) and normalized to the average ratio of all replicates for a given subject. The reference bar depicts the color-based gradient representing lower ratios (red) to higher ratios (green), with moderate ratios in the yellow range.

With insight into sPDGFRβ isoforms and their role in vascular developmental still emerging [45], we conducted a western blot analysis of murine placenta protein from each time point, labeling for PDGFRβ and α-tubulin as a loading control. Both full-length (∼160kDa) and soluble (∼32kDa) PDGFRβ protein increased from E9.5 to E14.5, with significant increases in both forms from E14.5 to E18.5 (Figure 3G-H), mirroring the transcriptional changes observed for both. To begin exploring the potential relevance of sPDGFRβ isoforms to human vascular development and maturation, we collected placenta protein samples from full-term, uncomplicated pregnancies, sampling from 4 spatially distinct quadrants. Western blot analysis of these samples revealed the presence of full-length PDGFRβ (∼160kDa) as well as a truncated PDGFRβ variant (∼55kDa) (Figure 3I). For each replicate, a ratio of integrated densities for PDGFRβ bands and the corresponding α-tubulin band was calculated and then normalized to the average for all replicates for a given subject. We used these values to generate a heat-map for relative PDGFRβ levels (Figure 3J), providing a visual representation of the range and regional heterogeneity in PDGFRβ protein within the human placenta.

To begin addressing sPDGFRβ variant regulation and function during placental vascular development, we sought a cell-based model that could capture key features of placenta vessel development including PC differentiation and sPDGFRβ production. We recently characterized murine wild-type (WT) and “double-reporter” (DR) embryonic stem cell (ESC) models that give rise to primitive vascular structures involving PCs upon spontaneous differentiation [68]. Follow-on analysis of these models suggested that TB stem cells (TSC) – a lineage giving rise to placenta villous cytotrophoblasts and intermediate trophoblasts [78] – was potentially present within these stem cell models also. We pursued several lines of evidence to determine the relative abundance of TSCs during stem cell differentiation. First, we immunostained E14.5 murine placenta sections for PECAM-1/CD31 to label ECs alongside staining for cytokeratin-8 (TROMA-I/*Krt8*) to verify this immunolabeling tool for detecting TB lineage cells [79] (Figure 4Ai-iii). Applying the same staining protocol to WT murine stem cells differentiated for 8 days revealed PECAM-1/CD31-positive vascular structures adjacent to numerous cells labeled for TROMA-I (Figure 4Aiv-vi). Villous cytotrophoblasts and syncytio-trophoblasts are also known to express the tight junction protein zonula occludens-1 (ZO-1/*Tjp1*) as they form important tissue barriers during placenta development [80]. We therefore immunostained for ZO-1 in DR murine stem cells at differentiation day 10 (dd10) when primitive vascular structures could be visualized via the *Flk-1:eGFP* reporter in the emerging endothelium (Figure 4Bi & iv) and *NG2:DsRed* in PCs and their progenitors (Figure 4Bii & iv). ZO-1 staining clearly labeled the cell-cell borders of TB lineage cells, while also appearing within a subset of ECs within maturing vessels (Figure 4Biii & iv). We corroborated these staining patterns with transcriptional data generated previously by RNA sequencing of unlabeled, flow-through cells (i.e. lacking eGFP or DsRed reporters) collected via FACS from undifferentiated, dd7, and dd10 DR stem cells [68]. In addition to increased expression of cytokeratin-8 (*Krt8*) and ZO-1 (*Tjp1*) over time, other commonly accepted TB lineage markers [81] were upregulated including *Krt18, Krt7, Cdh1, Rrm2, Hand1, Gata2,* and *Gata3* (Figure 4C). These gene expression changes coincided with robust vessel formation, which was visualized via DR labeling of emerging ECs and PC progenitors at dd8, dd10, and dd12 (Figure 4D).

**Figure 4.**
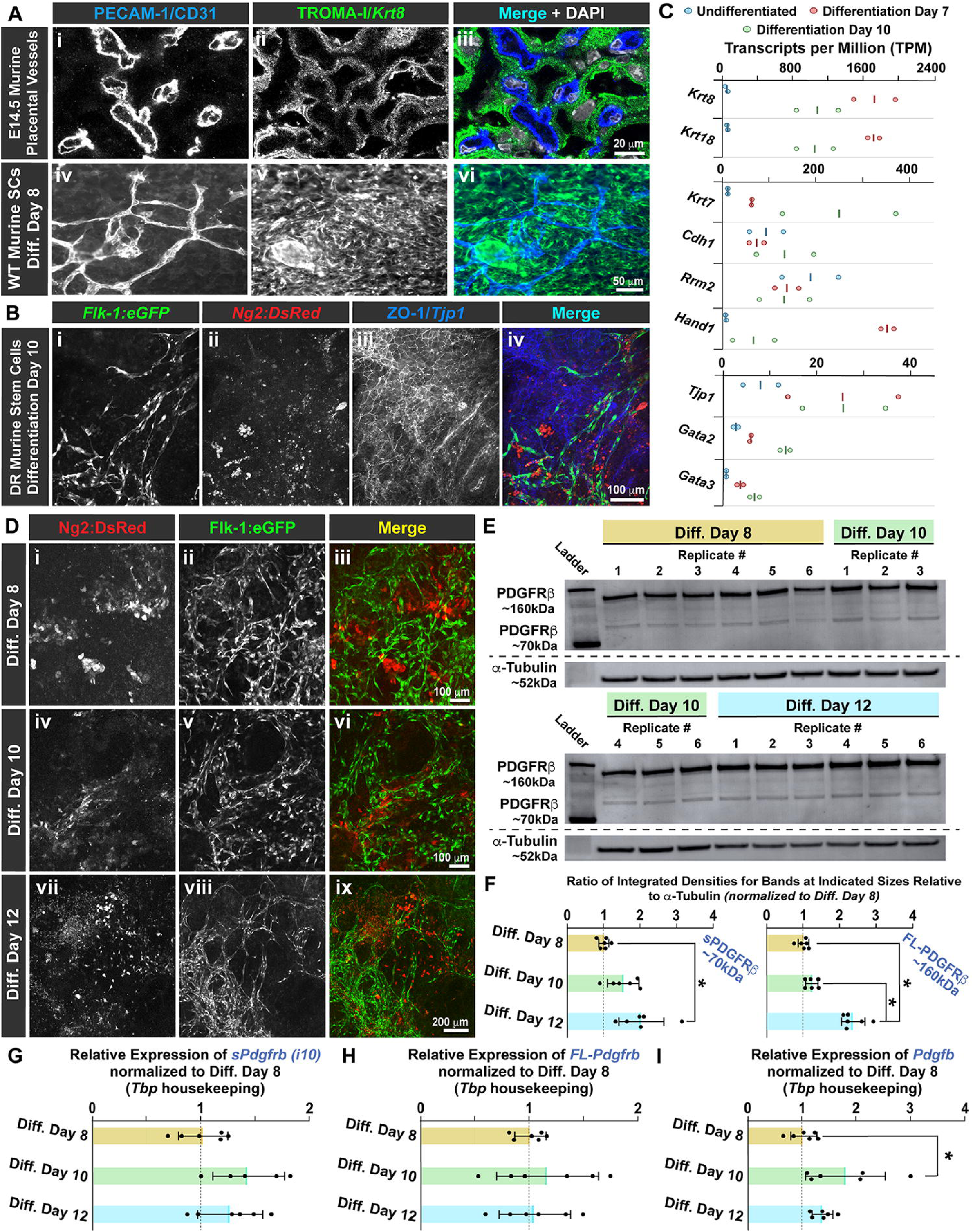
Differentiation of Murine Stem Cells Yields a Trophoblast-Like Lineage alongside the Formation of Primitive Vascular Structures involving Pericyte Precursors and Soluble PDGF Receptor-β Isoform Production. (A) Representative confocal images of an E14.5 murine placental section (i-iii) and WT murine stem cells at differentiation day 8 (dd8, iv--vi) demonstrating immunofluorescent labeling of endothelial cells via PECAM-1/CD31 antibody (i & iv; blue in iii & vi) and trophoblast-like cells via TROMA-I/Krt8 antibody (ii & v; green in iii & vi). Merged overlays include cell nuclei labeling by DAPI (white in iii & vi). Scale bars are 20 mm in iii, and 50 mm in vi. (B) Representative confocal images of “double reporter” (DR) murine stem cells at differentiation day 10 (dd10) demonstrating reporter and immunofluorescent labeling of endothelial cells via *Flk-1^eGFP/+^*reporter (i; green in iv), pericyte precursors via *NG2/Cspg4^DsRed/+^*reporter (ii; red in iv), and trophoblast-like cells via ZO-1/*Tjp1* antibody (iii; blue in iv). Merged overlay includes cell nuclei labeling by DAPI (white in iv). Scale bar is 100 mm. (C) Graphs of RNA sequencing measuring transcripts per million (TPM) for GFP- and DsRed-negative cells from the unsorted fraction following FACS of undifferentiated stem cells (blue ovals, n=2), those at differentiation day 7 (dd7, red ovals, n=2), and dd10 (green ovals, n=2). Gene targets include commonly accepted trophoblast lineage markers: *Krt8, Krt18, Krt7, Cdh1, Rrm2, Hand1, Tjp1, Gata2, & Gata3.* Lines indicate averages. (D) Representative images of DR murine stem cells differentiated for 8 (i-iii), 10 (iv-vi), and 12 days (vii-ix). Pericyte progenitors labeled with DsRed (*NG2/Cspg4^DsRed/+^*reporter in i, iv, & vii; red in iii, vi, & ix) emerged alongside the developing endothelium expressing eGFP (*Flk-1^eGFP/+^* reporter in ii, v, & viii; green in iii, vi, & ix) forming nascent vascular networks. Merged images show both channels (iii, vi, & ix), with respective scale bars: 100 mm in iii and vi, and 200 mm in ix. (E) Representative western blot images of protein from differentiated murine stem cells labeled for PDGFRβ (∼160kDa and ∼70kDa) with α-tubulin (∼52kDa) serving as loading control. *Top gel:* Ladder in Lane 1, 6 replicates from differentiation day 8 in Lanes 2-7, & 3 replicates from differentiation day 10 in Lanes 8-10. *Bottom gel:* Ladder in Lane 1, 3 replicates from differentiation day 10 in Lanes 2-4, & 6 replicates from differentiation day 12 in Lanes 5-10. *Images were cropped and inverted from original blots, which are provided in Supplemental* Figures 5-6. (F) Graph of ratios of integrated densities for bands at the indicated sizes (∼70kDa and ∼160kDa) relative to α-tubulin (loading control run on same blot). Individual data points are shown with averages represented as bars (dd8--tan, dd10--green, dd12--blue), with dotted gray lines indicating reference to dd8 samples. *p≤0.05 by one-way ANOVA with post-hoc Tukey test for comparisons indicated. *Note in (G-I), for each gene expression graph shown: all data were normalized to dd8 as indicated by the dotted line, Tbp served as the housekeeping gene, individual data points were plotted for each differentiation day with averages represented by bars: dd8--tan, dd10--green, dd12--blue, error bars represent standard error of the mean, and *p*≤*0.05 by one-way ANOVA with post-hoc Tukey test for comparisons indicated.* (G) Graph of relative *Pdgfb* expression within differentiating murine stem cells at dd8 (n=6), dd10 (n=6), and dd12 (n=6). (H) Graph of relative *FL-Pdgfrb* expression within differentiating murine stem cells at dd8 (n=6), dd10 (n=6), and dd12 (n=6). (I) Graph of relative *soluble Pdgfrb* expression (intron10 truncation variant) expression within differentiating murine stem cells at dd8 (n=6), dd10 (n=5), and dd12 (n=6).

Observing vascular development and TB lineage cells during differentiation within these murine stem cell models, we then wanted to determine if sPDGFRβ variants were also generated during early vessel formation. To do so, we collected cell lysates from DR stem cells at differentiation days 8, 10, and 12 and analyzed these proteins for the presence of sPDGFRβ isoforms by western blot. Along with bands at ∼160kDa corresponding to full length PDGFRβ, we found distinct bands at ∼70kDa labeled by the PDGFRβ antibody, aligning with the predicted size of sPDGFRβ variants generated by alternatively spliced transcripts truncated within intron 10 (Figure 4E). Assessing the integrated densities of these bands and normalizing them to the α-tubulin loading control bands, we found a steady increase in sPDGFRβ abundance over time, while full length PDGFRβ increased more distinctly at dd12 (Figure 4F). On the transcriptional level, the intron 10 (i10) sPDGFRβ variants displayed slight increases at dd10 and dd12, though expression remained steady over time, as did the levels of full length *Pdgfr*β (Figure 4G-H). *Pdgf-b* expression however increased significantly at dd10 relative to dd8 and decreased slightly at dd12 (Figure 4I). Taken together, these data demonstrate that murine stem cell differentiation can give rise to TB lineage cells and placenta-associated vascular structures including PCs, while also involving the production of sPDGFRβ isoforms.

### During Early Vessel Formation, Soluble PDGFR**β** Levels Appear to be Independent of ADAM10 Activity and Exogenous Growth Factors but Sensitive to Hypoxia, in Part to Regulate Full Length PDGFR**β** Dynamics

Observing sPDGFRβ isoform production alongside vessel formation and TB progenitors in our murine TESC differentiation model, we next wanted to utilize this model to begin identifying mechanisms that may contribute to sPDGFRβ regulation. Previous studies have suggested that the metalloprotease ADAM10 may cleave full length PDGFRβ to generate soluble fragments, though primarily in the context of PC damage and neurological pathology [44, 46, 47]. To test the hypothesis that ADAM10 may give rise to sPDGFRβ isoforms during early vascular development, we applied the ADAM10 inhibitor GI-254023X [82] or DMSO vehicle control to differentiating TESCs at dd8 and dd10, collecting protein lysates on dd12 for western blot analysis (Figure 5A). ADAM10 inhibition had no effect on sPDGFRβ (∼70kDa) protein levels and caused a significant increase in full length PDGFRβ (∼160kDa) (Figure 5B-C), contrary to our initial expectations. These data suggest that sPDGFRβ production may not depend on ADAM10 activity in the context of placental vascular development, perhaps relying more on mRNA alternative splicing.

**Figure 5.**
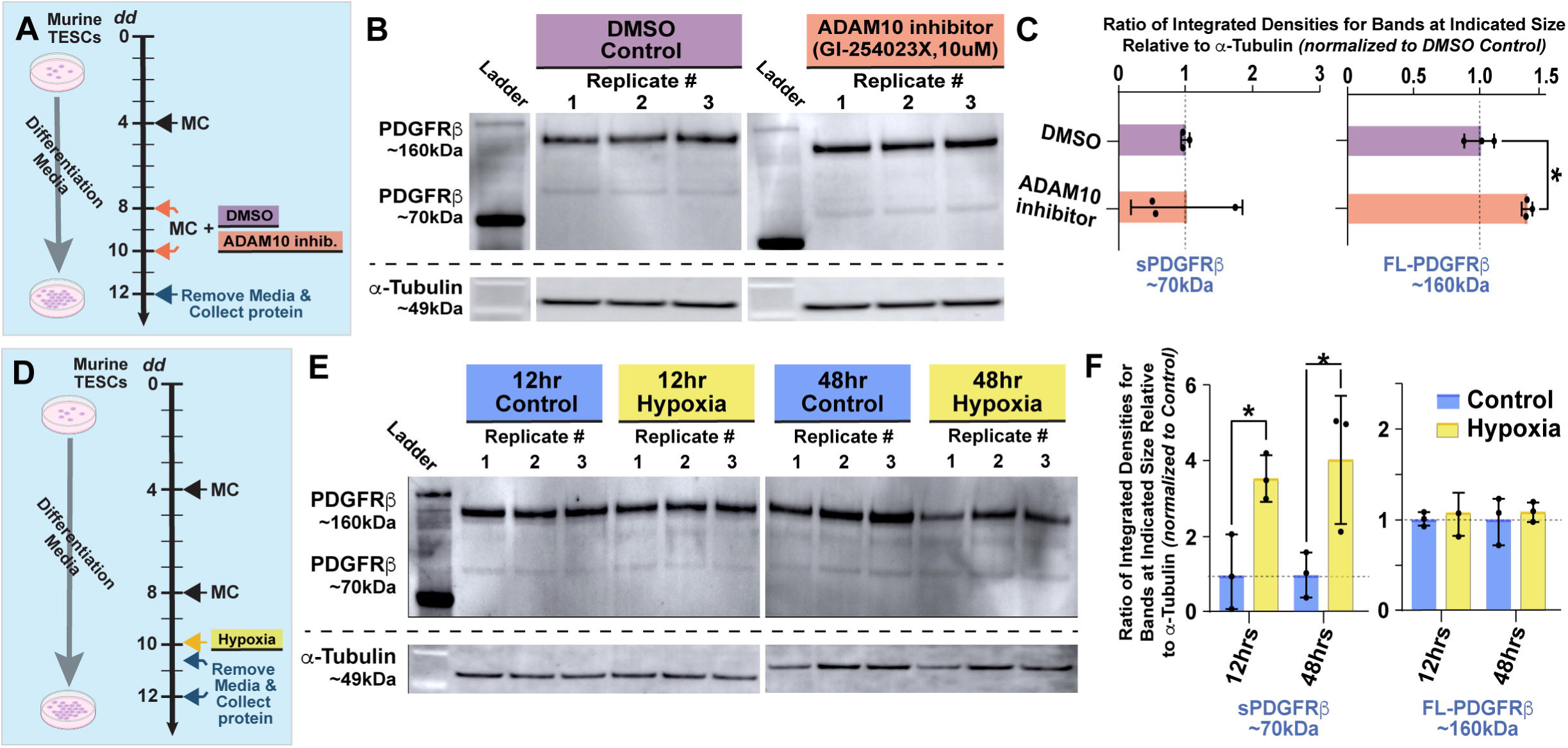
ADAM10 Inhibition Does Not Affect Soluble PDGF Receptor-β Levels during Murine Stem Cell Differentiation into Primitive Vessels, while Hypoxia Increases Isoform Levels. (A) Schematic of murine trophoblast and embryonic stem cell (TESC) differentiation experiment over 12 days. Media Changes (MCs) occurred at differentiation day (dd) 4 and 8, with dd8 including treatment with DMSO (vehicle control) or the ADAM10 inhibitor GI-254023X in DMSO (10uM). At dd12, media was removed, cells washed with PBS, and cell protein collected for western blot. (B) Representative western blot images of protein from differentiated murine stem cells labeled for PDGFRβ (∼160kDa and ∼70kDa) with α-tubulin (∼52kDa) serving as loading control. Ladder in Lane 1 of Blot #1 and 2, 3 replicates from DMSO control group in Lanes 2-4 of Blot #1, & 3 replicates from ADAM10 inhibitor treatment group in Lanes 2-4 of Blot #2. *Images were cropped and inverted from original blots, which are provided in Supplemental* Figures 7-8. (C) Graph of ratios of integrated densities for bands at the indicated sizes (∼70kDa and ∼160kDa) relative to α-tubulin (loading control run on same blot). Individual data points are shown with averages represented as bars (DMSO control—purple, n=3, ADAM10 inhibitor—pink, n=3), with dotted gray lines indicating reference to DMSO control samples and error bars representing standard error of the mean. *p≤0.05 by one-way ANOVA with post-hoc Tukey test for comparisons indicated. (D) Schematic of murine trophoblast and embryonic stem cell (TESC) differentiation experiment over 12 days. Media Changes (MCs) occurred at differentiation day (dd) 4 and 8, with dd10 being on the onset of hypoxia (3% O_2_) or remaining under normal culture conditions (control). At dd10.5 (12hr duration) or dd12 (48hr duration), media was removed, cells washed with PBS, and cell protein collected for western blot. (E) Representative western blot images of protein from differentiated murine stem cells labeled for PDGFRβ (∼160kDa and ∼70kDa) with α-tubulin (∼52kDa) serving as loading control. Ladder in Lane 1 of Blot #1, 3 replicates from 12hr control group in Lanes 2-4 of Blot #1, 3 replicates from 12hr hypoxia group in Lanes 5-7 of Blot #1, 3 replicates from 48hr control group in Lanes 1-3 of Blot #2, & 3 replicates from 48hr hypoxia group in Lanes 4-6 of Blot #2. *Images were cropped and inverted from original blots, which are provided in Supplemental Figure 9-10.* (F) Graph of ratios of integrated densities for bands at the indicated sizes (∼70kDa and ∼160kDa) relative to α-tubulin (loading control run on same blot). Individual data points are shown with averages represented as bars (control—blue, n=3 for each time point, 3% O_2_ hypoxia—yellow, n=3 for each time point), with dotted gray lines indicating reference to control samples and error bars representing standard error of the mean. *p≤0.05 by one-way ANOVA with post-hoc Tukey test for comparisons indicated. Schematics incorporated elements from Biorender.com.

Hypoxia has also been shown to affect sPDGFRβ protein levels in several cell-based models [44, 45]. Here, we subjected differentiating TESCs to 3% O_2_ for 12hrs and 48hrs, with control samples remaining at standard cell culture oxygen tension (∼21% O_2_) over the same time points (Figure 5D). Western blot analysis of protein lysates revealed significant increases in sPDGFRβ (∼70kDa) protein levels at 12hrs and 48hrs of reduced oxygen tension, while full length PDGFRβ (∼160kDa) remained unchanged over these durations of relative hypoxia (Figure 5E-F). These experiments indicate that mechanisms governing sPDGFRβ generation are likely oxygen sensitive.

Several growth factor pathways, particularly within the RTK family, are known to include “decoy” receptors as a feedback mechanism to modulate signaling, e.g. Flt-1 [57–59]. The relative levels of growth factor ligand present can often influence these mechanisms [50–55]. We hypothesized therefore that exogenous growth factor exposure might affect sPDGFRβ regulation, perhaps causing an increase in isoform levels. To test this idea, we differentiated TESCs under standard conditions for 8 days and then transitioned cultures to control or experimental media conditions (Figure 6A). Specifically, control samples received an endothelial growth media (EGM) with reduced fetal bovine serum (FBS) levels (10%) and lacking additional growth factors or supplements. One experimental group was cultured in EGM with 10% serum and added VEGF-A, with another group exposed to EGM plus 10% serum and added PDGF-BB. The final experimental group involved the complete EGM kit provided by Lonza^TM^ with additional PDGF-BB, i.e. EGM + 10% FBS + VEGF-A + insulin growth factor-2 (IGF-2) + EGF + PDGF-BB + ascorbic acid + heparin. These media configurations were changed on dd10, and protein lysates were collected on dd12 following media removal and PBS wash. Western blot analysis of these proteins revealed lower levels of sPDGFRβ isoforms (∼70kDa) with each of the different media configurations relative to control conditions, though no reductions were statistically significant (Figure6B-C). Full length PDGFRβ (∼160kDa) levels were unaffected by VEGF-A exposure; however, the inclusion of PDGF-BB with the basic EGM or the EGM complete kit caused a significant reduction in FL-PDGFRβ protein levels, consistent with ligand binding induction of receptor internalization and degradation [64].

**Figure 6.**
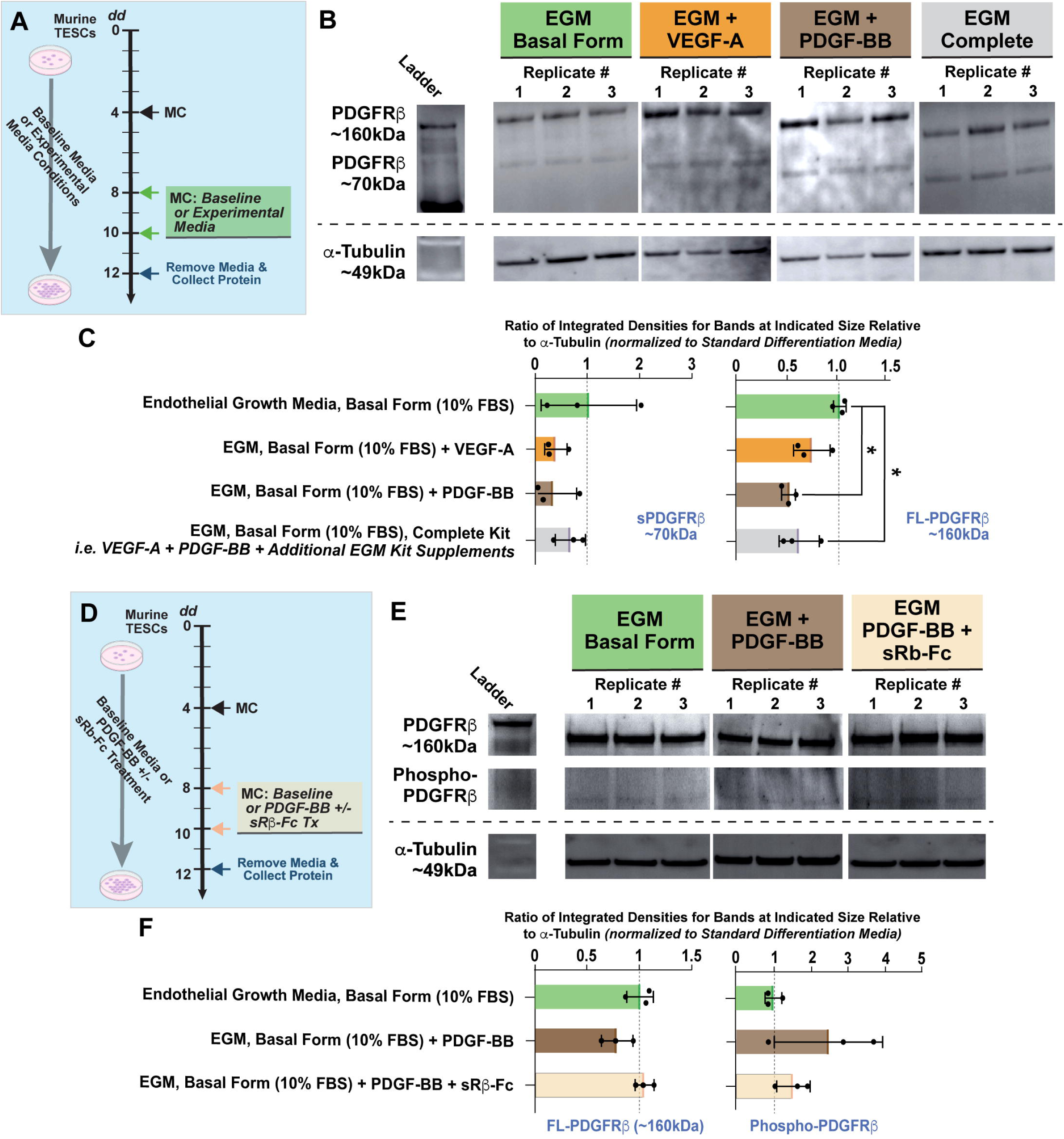
Exogenous Growth Factors Decrease Full Length PDGFRβ Protein Levels during Formation of Murine Stem Cell-Derived Vessels, with sPDGFRβ Levels Remaining Constant, but Adding an sPDGFRβ Mimetic Partially Rescues Growth Factor-Induced Changes in Full Length PDGFRβ Levels and Phosphorylation. (A) Schematic of murine trophoblast and embryonic stem cell (TESC) differentiation experiment over 12 days. Media Changes (MCs) occurred at differentiation day (dd) 4, 8, and 10 with dd8 and dd10 involving transition to Endothelial Growth Media (EGM), Basal Form (10% FBS) alone, with added VEGF-A, added PDGF-BB, or addition of the Complete EGM Kit. At dd12, media was removed, cells washed with PBS, and cell protein collected for western blot. (B) Representative western blot images of protein from differentiated murine SCs labeled for PDGFRβ (∼160kDa and ∼70kDa) with α-tubulin (∼52kDa) serving as loading control. Ladder in Lane 1 (from Lane 1, Blot #2), 3 replicates from EGM Basal Form control group in Lanes 2-4 (from Lanes 6-8, Blot #2), 3 replicates from EGM Basal Form + VEGF-A group in Lanes 5-7 (from Lanes 8-10, Blot #1), 3 replicates from EGM Basal Form + PDGF-BB group in Lanes 8-10 (from Lanes 5-7, Blot #1), & 3 replicates from EGM Basal Form + Complete EGM Kit group in Lanes 11-13 (from Lanes 3-5, Blot #2). *Images were cropped and inverted from original blots, which are provided in Supplemental Figure 11-12.* (C) Graph of ratios of integrated densities for bands at the indicated sizes (∼70kDa and ∼160kDa) relative to α-tubulin (loading control run on same blot). Individual data points are shown with averages represented as bars (EGM Basal Form control—green, n=3; EGM Basal Form + VEGF-A group—orange, n=3; EGM Basal Form + PDGF-BB group—brown, n=3; EGM Basal Form + Complete EGM Kit group—gray, n=3), with dotted gray lines indicating reference to EGM Basal Form control samples and error bars representing standard error of the mean. *p≤0.05 by one-way ANOVA with post-hoc Tukey test for comparisons indicated. (D) Schematic of murine TESC differentiation experiment over 12 days. MCs occurred at dd4, 8, and 10 with dd8 and dd10 involving transition to Endothelial Growth Media (EGM), Basal Form (10% FBS) alone, with added PDGF-BB, or with the addition of PDGF-BB and sRβ mimetic. At dd12, media was removed, cells washed with PBS, and cell protein collected for western blot. (E) Representative western blot images of protein from differentiated murine SCs labeled for PDGFRβ (∼160kDa and ∼70kDa) with α-tubulin (∼52kDa) serving as loading control. Ladder in Lane 1 (from Lane 1, Blot #1), 3 replicates from EGM Basal Form control group in Lanes 2-4 (from Lanes 5-7, Blot #1), 3 replicates from EGM Basal Form + PDGF-BB group in Lanes 5-7 (from Lanes 8-10, Blot #2), & 3 replicates from EGM Basal Form + PDGF-BB and sRβ-Fc mimetic group in Lanes 8-10 (from Lanes 2-4, Blot #1). *Images were cropped and inverted from original blots, which are provided in Supplemental Figure 13-14.* (F) Graph of ratios of integrated densities for bands at the indicated sizes (∼70kDa and ∼160kDa) relative to α-tubulin (loading control run on same blot). Individual data points are shown with averages represented as bars (EGM Basal Form control—green, n=3; EGM Basal Form + PDGF-BB group—brown, n=3; EGM Basal Form + PDGF-BB and sRβ-Fc mimetic group—tan, n=3), with dotted gray lines indicating reference to EGM Basal Form control samples and error bars representing standard error of the mean. Schematics incorporated elements from Biorender.com.

Having observed this reduction in FL-PDGFRβ levels with PDGF-BB treatment, we next wanted to ask if co-administration of an sPDGFRβ mimetic could rescue this phenomenon and provide insight into potential sPDGFRβ function. To do this, we utilized a commercially available peptide containing the PDGF-BB ligand binding domain, mirroring the same region present in sPDGFRβ isoforms [45]. This mimetic peptide, which also harbors a human Fc receptor region that enhances protein stability, has been previously shown to modulate PDGF-BB signaling and PC behaviors [33]. We utilized the same experimental approach as in the exogenous growth factors experiment with a control group of EGM plus 10% serum alone and an experimental group involving EGM with 10% FBS and added PDGF-BB, both beginning at dd8 (Figure 6D). Here, we also included a group exposed to EGM + 10% FBS + PDGF-BB + sPDGFRβ-Fc mimetic (sRβ-Fc). Protein lysates were collected at dd12 and analyzed for FL-PDGFRβ levels alongside the relative amount of FL-PDGFRβ phosphorylation. As in the exogenous growth factor experiment, we found that PDGF-BB treatment alone caused a reduction in FL-PDGFRβ levels; the sPDGFRβ mimetic however restored those levels back to control values (Figure 6E-F). Full length PDGFRβ phosphorylation demonstrated a corresponding increase with PDGF-BB treatment, and the addition of the sPDGFRβ mimetic dampened that increase back towards control levels of PDGFRβ phosphorylation. Overall, these data suggest that, in our TESC model of early vessel formation, sPDGFRβ protein levels do not depend on ADAM10 activity or excess growth factor levels, but they increase during acute hypoxia and can potentially modulate full length PDGFRβ phosphorylation and associated internalization and degradation dynamics.

## DISCUSSION

Vascular remodeling during fetal and placenta formation is critical for sustaining the growth and health of all developing tissues and organs. Capillary PCs in particular promote the stability and maturity of microvascular networks by extending along ECs and contributing to vessel integrity, among other functions [22, 83, 84]. Pericyte-EC crosstalk in the placenta, as in all tissues, is facilitated in part by the PDGF-BB pathway [18, 43]. This key signaling axis orchestrates PC differentiation, proliferation, migration, and survival, culminating in PC recruitment and retention within the capillary wall [28, 30]. Here, we found morphological and molecular evidence in murine and human placenta tissues for progressive PC investment along feto-placental capillaries during gestation. Placental PCs also exhibited a phenotypic heterogeneity as seen in other tissues such as the brain, heart, and lung [60–62]. Alongside the molecular mechanisms coordinating vessel formation and PC expansion, we found that the recently identified soluble isoforms of PDGFRβ [45] were present in murine and human placentas on the protein and mRNA transcript levels. Soluble PDGFRβ variant expression was also found in a model of murine stem cell differentiation [68] that recapitulated key aspects of early placental vascular development, i.e. blood vessel formation and remodeling involving PCs, and the emergence of TB lineage cells. Using this TESC differentiation model, we found that sPDGFRβ levels were not dependent on ADAM10 activity or exogenous growth factors during early vascular development. Hypoxia however caused a dramatic increase in sPDGFRβ isoforms detected. To better understand how elevated levels of sPDGFRβ variants might affect PDGF-BB signaling, we exposed TESC-derived vessels to excess PDGF-BB with and without a sPDGFRβ mimetic. PDGF-BB treatment alone caused a reduction in full length PDGFRβ with receptor phosphorylation being increased, consistent with known regulatory mechanisms [64]. The inclusion of the sPDGFRβ mimetic alongside PDGF-BB treatment shifted these changes back toward control levels, consistent with a potential feedback role observed in other studies [33, 65, 66]. Overall, these data suggest that, during murine and human placenta development, PCs are recruited to capillaries at the maternal-fetal interface, exhibiting a phenotypic heterogeneity comparable with other organs. Our data further suggest that soluble PDGFRβ isoforms may influence full length PDGFRβ signaling dynamics in this context, likely independent of ADAM10 activity and growth factor levels but potentially sensitive to reduced oxygen tension.

Pericytes are a nearly ubiquitous component of the capillary wall across many tissue beds and organs. As such, they have been assigned a wide array of functions including as tissue resident stem cells and in endothelial barrier reinforcement, as angiogenic stimuli and modulators of capillary tone, among many other disparate roles [22, 83, 84]. While their global role across all tissues remains ill-defined, it is becoming clearer that PC subtypes exist. We and others have found that PC subpopulations exhibit distinct morphological and transcriptional characteristics [60–62, 70]. Pericyte phenotypic heterogeneity is an emerging concept in the placenta as well, though recent studies have begun shedding light on this concept [2, 14]. In the current study, we found that placental PCs appear to express many of the common markers found in embryonic and adult tissues such as NG2, PDGFRβ, Notch3, and Vitronectin [25, 26, 68, 74–76, 83]. Interestingly, we found NG2-positive PCs adjacent to capillary-sized vessels with heterogeneous expression of the contractility protein αSMA, suggesting subsets of placenta PCs may have differential levels of contractility. Where PCs have been described as contractile such as in the brain, they have been suggested to respond rapidly to regionally heterogeneous fluctuations in metabolic demand, such as in response to neuronal firing [85, 86]. While this is an intriguing interpretation of PC function, it is not clear how this role would apply to the placenta, which lacks innervation [87] and is unlikely to require that same level of fine-grain control over feto-placental capillary perfusion. Thus, observing a phenotypic heterogeneity in placental PC populations (here and in [2, 14]) that has been described in other tissues and organs [60–62] raises the question of their universal role within the microcirculation, especially with a mix of contractile and seemingly non-contractile PC subtypes. Follow-on studies will be needed to further resolve the necessity for PC diversification in supporting the function of capillary beds across all tissues and organs where they are found, especially the placenta.

Pericytes have been implicated in processes related to angiogenic remodeling as well as to capillary maturation and transition towards quiescence. As such, their involvement in vascular development is often positioned towards the latter stages of vessel formation. Recent studies from our lab have suggested that PCs and/or their precursors are present and engaging the developing endothelium at earlier stages of embryonic development including vasculogenesis [68, 84]. Here, we collected additional in vivo evidence supporting an alternative timeline for their involvement as PC lineage cells within the placenta were found alongside vasculogenic endothelium in the placenta that seemed to harbor regions described as hemogenic [88]. This timeline for vascular cell differentiation and morphological organization was reflected in our stem cell model as well. We further found that, during stem cell differentiation, TB lineage cells emerged. During the generation of embryonic stem cell lines, it is often difficult to avoid the inclusion of the trophectoderm, i.e. the cells constituting the outermost layer of the mammalian blastocyst that facilitate implantation and contribute to placenta formation [89]. In the current study, we asked if PC lineage cells and vasculogenic endothelium were emerging in conjunction with TB differentiation. Indeed, we found evidence for vascular development occurring adjacent to early placentogenesis in our stem cell differentiation model. Trophoblast lineage cells are known to be enriched for cytokeratins [81]; here, we detected increases in cytokeratin-7, -8, and -18 transcripts by bulk RNA sequencing of non-vascular cells from our TESC model. We also observed cytokeratin-8 (TROMA-I) [79] by immunohistochemistry of differentiated stem cells, with antibody validation in E14.5 murine placenta tissue. Moreover, we found the tight junction accessory protein zonula occludens-1 (ZO-1/*Tjp1*) [80] upregulated on the transcriptional level in differentiating non-vascular cells. ZO-1 protein was observed at the cell-cell interface of the nascent endothelium, as expected [90], but we also found junction-associated signals in TB progenitors in differentiated TESCs. Overall, our data support revisiting the working timeline for PCs and their progenitors being involved in vascular development, particularly during placentogenesis, which appears to be reasonably approximated by TESC differentiation models, including the one presented herein.

Signaling pathways that govern PC investment and expansion appear to be conserved across placental and embryonic development. In particular, the PDGF-BB pathway is critical for orchestrating PC contributions to placenta vasculature, as genetic loss of this key regulator in murine models leads to a range of placenta abnormalities, among other defects [18, 43]. Observations from the current study reinforce that notion, further shedding light on the existence of soluble PDGFRβ isoforms that emerge under physiological conditions in the mouse and human placenta. Previous studies have suggested the enzymatic cleavage by ADAM10 is required for generating sPDGFRβ variants [44, 46, 47], but here we found that ADAM10 activity is likely not involved in sPDGFRβ production in a stem cell-based model of the developing placenta. Additionally, we found that sPDGFRβ levels decreased with exogenous growth factor treatment of stem cell-derived vessels, though control groups suffered from high levels of variability that limited detection of statistically significant differences. Because angiogenic growth factor production is known to increase during hypoxia [91–94], we predicted that a reduction in oxygen tension might also lead to less sPDGFRβ generation in the stem cell model of placental vessel formation. We found however a robust and sustained increase in sPDGFRβ levels with 12- and 48-hours of hypoxia exposure, suggesting sPDGFRβ regulation might be independent of growth factor changes and may be impacted more by other regulatory mechanisms such as those related to mRNA alternative splicing [50–55]. Follow-on studies are aimed at this clarifying this question of sPDGFRβ regulation, specifically on contributions of enzymatic cleavage and the role of mRNA binding proteins [56] and splicing factors that give rise to transcript heterogeneity in the placenta and more broadly.

Soluble receptor isoforms have been implicated in a variety of key signaling pathways. The EGF pathway for example contains receptors that undergo ectodomain shedding of a soluble fragment that can influence full-length receptor signaling [48, 49]. This pathway, along with other RTK family members such as the FGF and VEGF-A pathways, include soluble receptors generated by alternatively spliced mRNA transcripts [50–55] with a level of regulation by RNA binding proteins [56]. Soluble Flt-1 (sFlt-1, or soluble VEGF Receptor-1 in humans) is a prime example of a truncated variant generated by mRNA alternative splicing that modulates VEGF-A signaling as a “decoy” receptor or ligand sink [57–59]. A close relative to VEGF within the RTK family, the PDGF pathway also appears to involve soluble isoforms of the full-length PDGFRβ signaling receptor, produced by truncated mRNA transcripts and perhaps enzymatic cleavage, though in specific contexts [44, 46, 47]. Cleaved sPDGFRβ is proposed to be a degradation product indicative of PC dysfunction or demise, with no specific or designated function. In the current study and in previous reports [45], however, sPDGFRβ mimetic peptides containing the ligand binding domain for PDGF-BB appear to modulate cell signaling and proliferation [65, 66] including disrupting PC expansion and interaction with remodeling ECs [33]. A subset of endogenously produced sPDGFRβ isoforms also appear to have heparin sulfate binding domains [45], suggesting potential interactions with the extracellular matrix that may influence its mode of action [95, 96]. Nevertheless, future work will be needed to more fully resolve its mechanistic involvement in maintaining PDGF-BB signaling within a specific range to achieve an appropriate level of PC association with capillaries during network remodeling.

While additional studies will be needed to establish a more complete picture of the potential clinical relevance for sPDGFRβ regulation and function, these isoforms and their associated transcripts are present in human cells and fluids under normal and pathological conditions. We are continuing efforts to validate sPDGFRβ mRNA transcript sequence information as well as amino acid sequence and protein structure, with the goal of using existing databases to screen for disease connections, particularly related to placental dysfunction. Given that soluble Flt-1 mis-regulation occurs during preeclampsia [6–8], we are eager to test the hypothesis that sPDGFRβ might also experience a level of aberrant regulation that may disrupt PC investment at the maternal-fetal interface. Furthermore, there is growing interest in how altered placentogenesis might impact fetal development, such as aberrant release of pro- and/or anti-angiogenic factors from the placenta that could disrupt fetal vascular development in key organs including the brain and heart [14, 15, 97]. Such a scenario could lead to vascular malformations in these and other organ systems, posing risk for life-long complications for the developing fetus as well as for acute pathologies e.g. unstable cerebrovasculature prone to rupture and giving rise to intraventricular hemorrhages [98, 99]. Thus, a translational approach will be needed to connect key molecular and cellular insights to clinical settings where sPDGFRβ might be used as a biomarker to stratify patients across disease risk severity and perhaps even serve as a viable therapeutic target for clinical management of vascular-related disease conditions.

## Supplemental Information

Supplemental information can be found online and includes supporting data from confocal image analysis using the machine learning-based platform QuPath along with original and unmarked western blots.

## Acknowledgements

We offer our sincerest thanks to Dr. Brittany R. Howell (Virginia Tech) and Dr. Kimberly P. Simcox (Carilion Clinic) for providing clinical samples. We would also like to thank all the members of the Chappell lab for their support and valued assistance on this project, both materially and intellectually. This work was supported in part by the Seale Foundation.

## Author Contributions

A.B., E.D., C.L., and J.C.C. conceptualized and optimized methodologies. A.B., E.D., C.L., and C.C. conducted experiments. A.B., E.D., R.D., C.L., C.C., C.S., and S.T. performed sample and dataset analysis. A.B. and J.C.C. wrote the manuscript. M.W., A.D., and J.C.C. obtained compliance approvals. B.H. and J.C.C. acquired financial support. All authors have reviewed and approved the final version of this manuscript.

## Data Availability

The datasets generated during the current study are included in the manuscript as individual data points where possible, and all supporting data are available from the corresponding author upon reasonable request.

## Abbreviations

sVEGFR1: soluble vascular endothelial growth factor receptor-1
PC: pericyte
EC: endothelial cell
TB: trophoblast
PDGF-BB: platelet-derived growth factor-BB
PDGF receptor-β: PDGFRβ-(or full length PDGFRβ -- FL-PDGFRβ)
sPDGFRβ: soluble PDGFRβ
EGF: epidermal growth factor
RTK: receptor tyrosine kinase
FGF: fibroblast growth factor
ADAM10: a disintegrin and metalloproteinase domain-containing protein 10
TESC: trophoblast and embryonic stem cell
PECAM-1: platelet-endothelial cell adhesion molecule-1
NG2: neural glial antigen-2
αSMA: α-smooth muscle actin
PlGF: Placental Growth Factor
DR: double reporter
ZO-1: zonula occludens-1
E#: embryonic day #
dd#: differentiation day #
EGM: endothelial growth media
IGF2: insulin growth factor-2

