## Supplemental Information for "Regulation of PDGF-BB Signaling in Placental Pericytes by Soluble PDGFRβ Isoforms: Implications for Fetoplacental Vascular Development"

**Supplemental Figure 1. Murine Chorio-allantoic Vasculature Progressively Expands during Gestation, with the Ratio of Pericytes to Endothelial Cells Remaining Relatively Constant.** (A) Graph of PECAM-1-positive cells normalized to all DAPI-positive cells for a given image from the indicated time points, E11.5 (n=3, green bar), E14.5 (n=3, orange bar), and E18.5 (n=4, purple bar). Individual data points shown, and errors bars are standard deviation. (B) Graph of the ratio between NG2-positive cells and PECAM-1-positive cells for a given image from the indicated time points, E11.5 (n=3, green bar), E14.5 (n=3, orange bar), and E18.5 (n=4, purple bar). Individual data points shown, and errors bars are standard deviation.

**Supplemental Figure 2.** Original western blots in support of Figure 3G.

**Supplemental Figure 3.** Original western blots in support of Figure 3I, Baby 001 and 002.

**Supplemental Figure 4.** Original western blots in support of Figure 3I, Baby 003 and 004.

**Supplemental Figure 5.** Original western blots in support of Figure 4E (Differentiation Day 8 and half of Differentiation Day 10 samples).

**Supplemental Figure 6.** Original western blots in support of Figure 4E (Half of Differentiation Day 10 and Differentiation Day 12 samples).

**Supplemental Figure 7.** Original western blots in support of Figure 5B.

**Supplemental Figure 8.** Original western blots in support of Figure 5B.

**Supplemental Figure 9.** Original western blots in support of Figure 5E.

**Supplemental Figure 10.** Original western blots in support of Figure 5E.

**Supplemental Figure 11.** Original western blots in support of Figure 6B.

**Supplemental Figure 12.** Original western blots in support of Figure 6B.

**Supplemental Figure 13.** Original western blots in support of Figure 6E.

**Supplemental Figure 14.** Original western blots in support of Figure 6E.

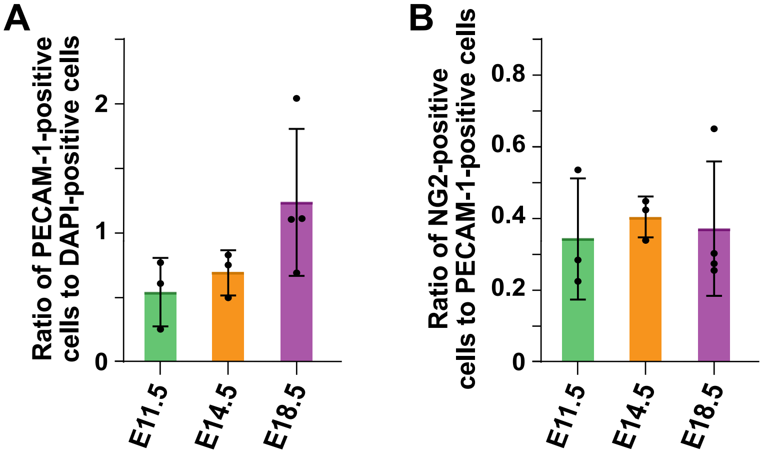


**Supplemental Figure 1.**

A

B

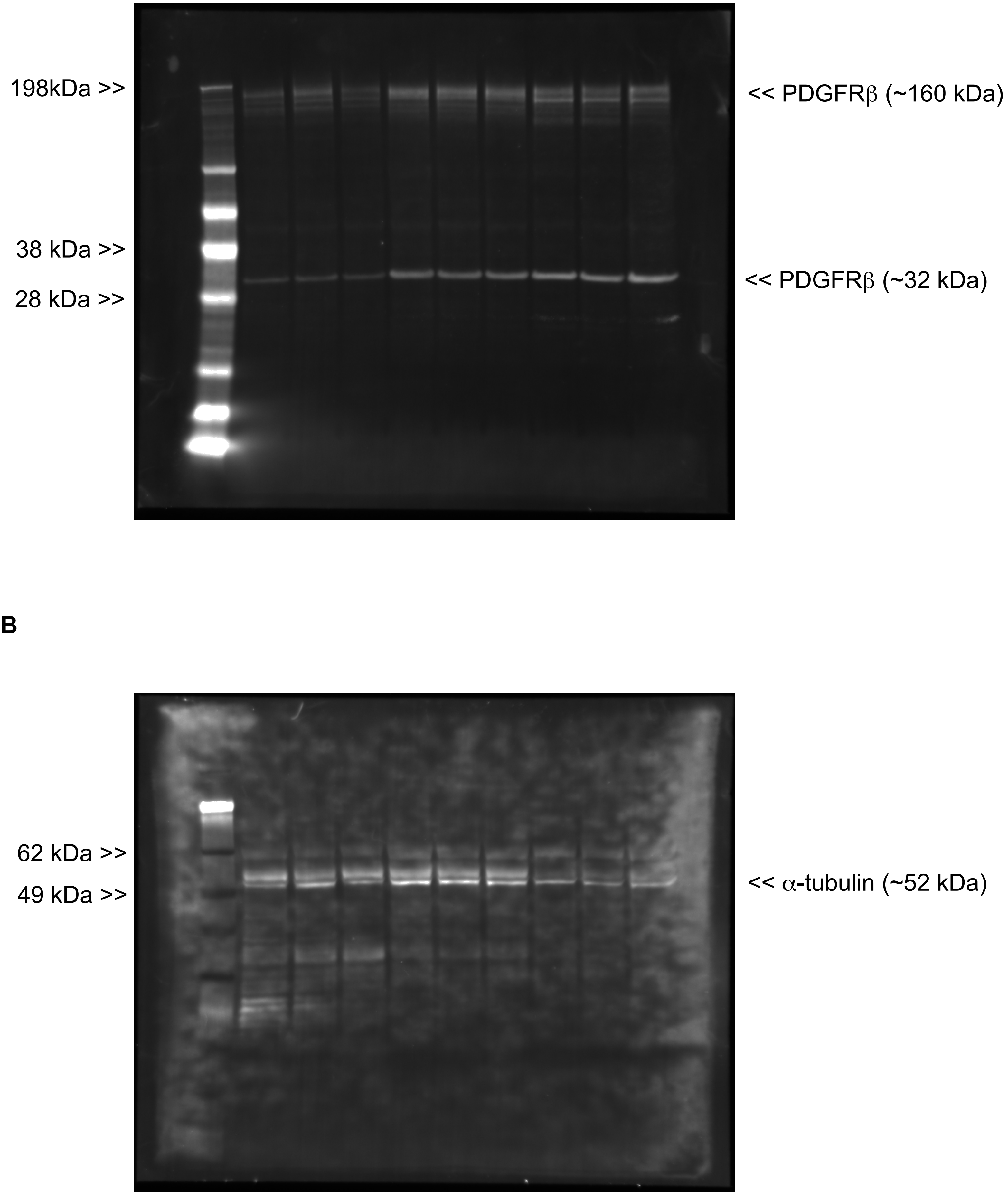


**Supplemental Figure 2.**

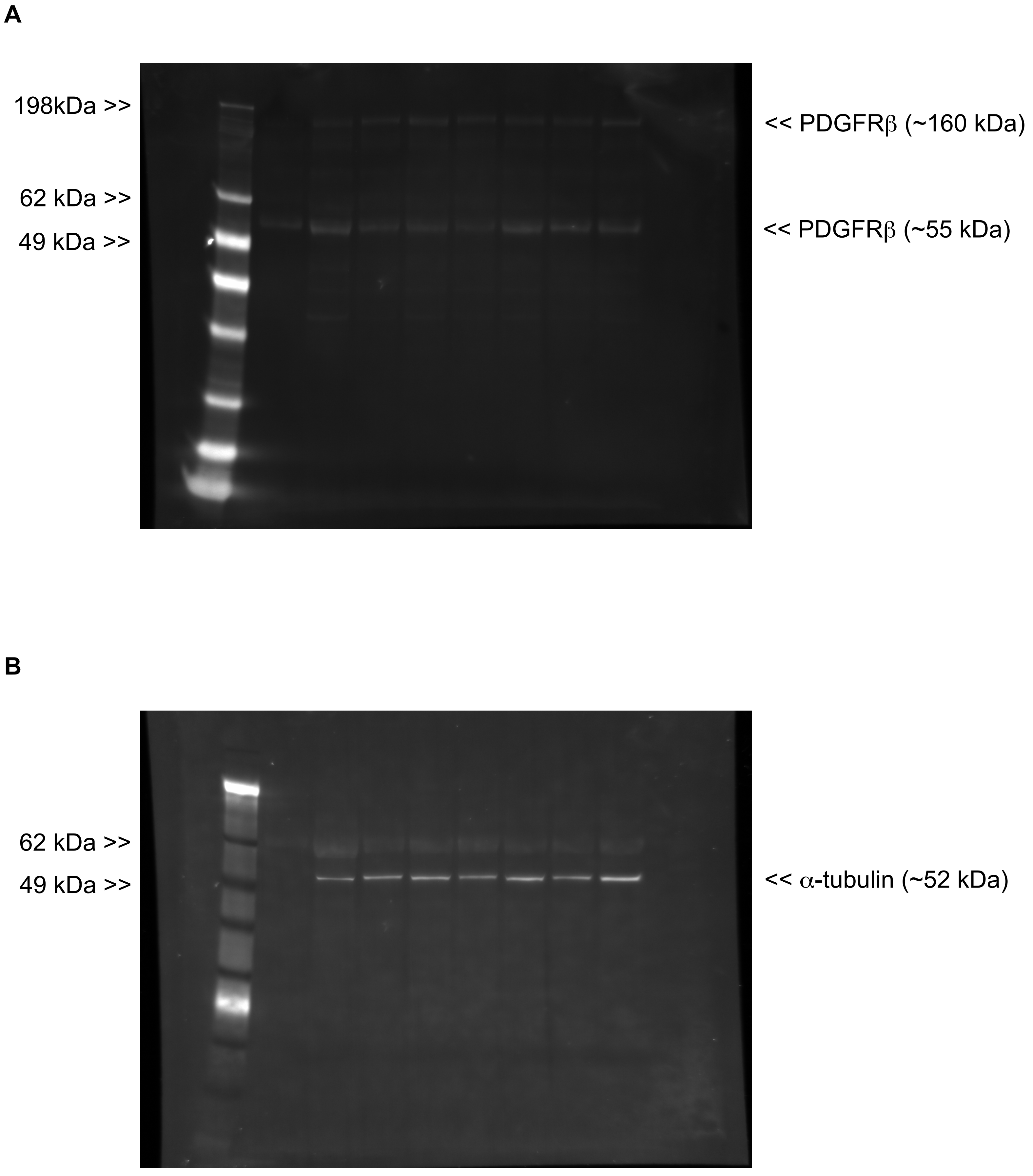


**Supplemental Figure 3.**

**A**

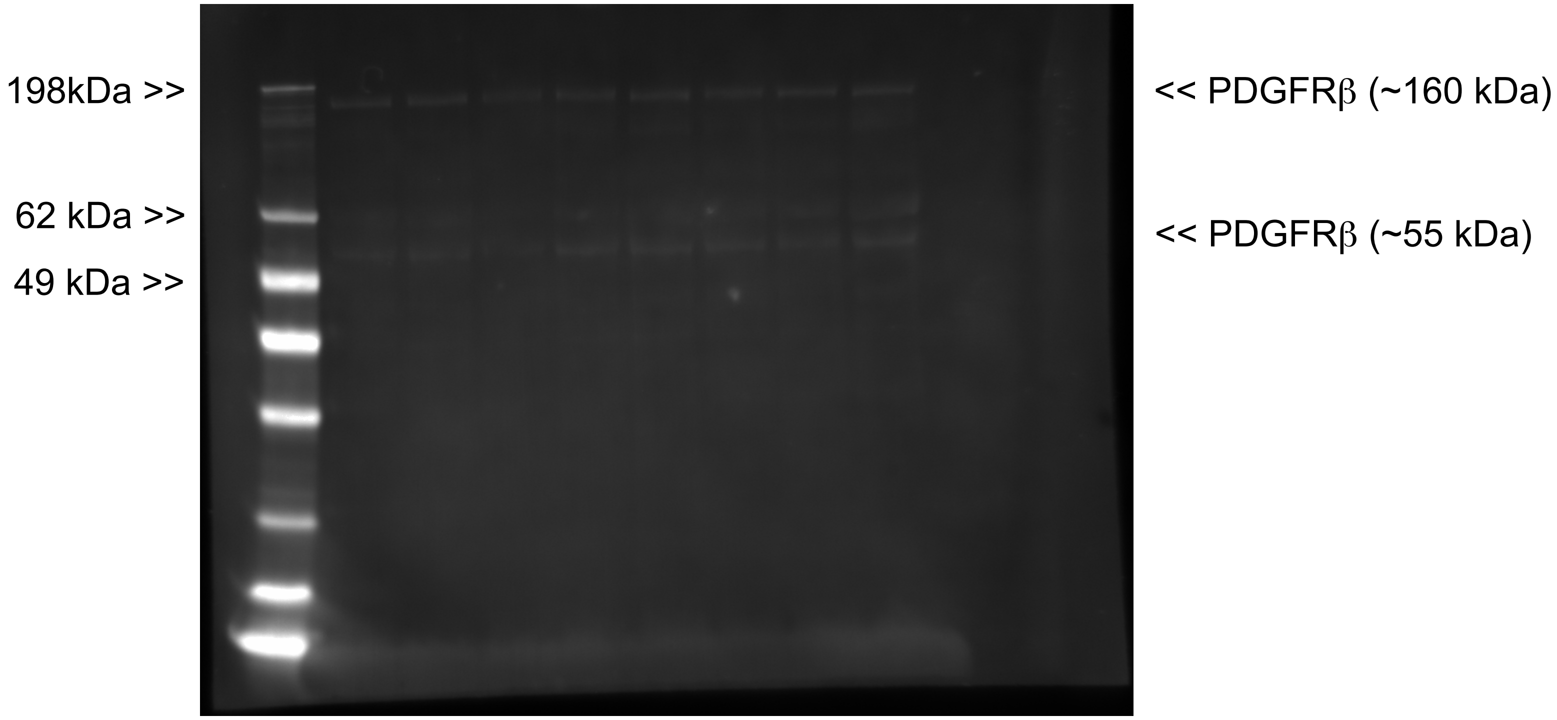


**B**

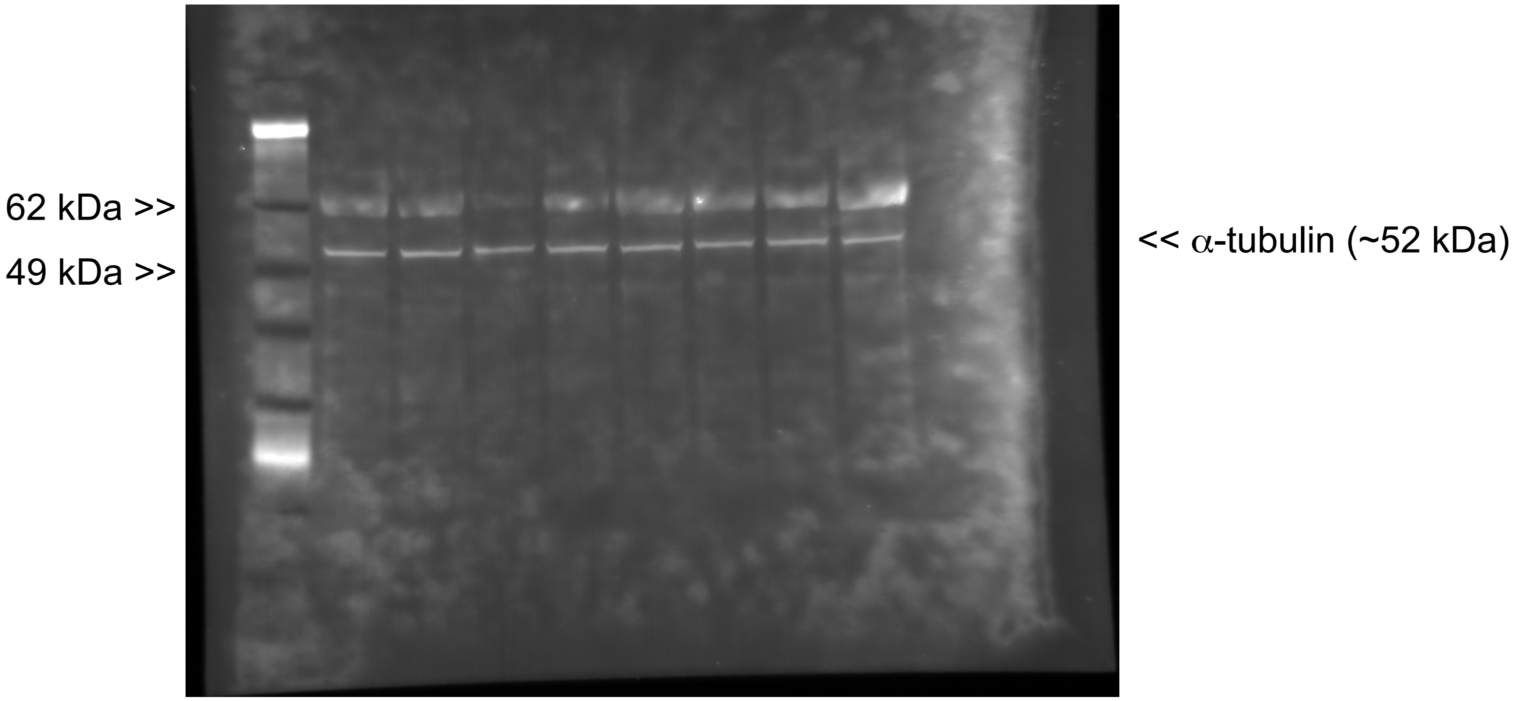


**Supplemental Figure 4.**

A

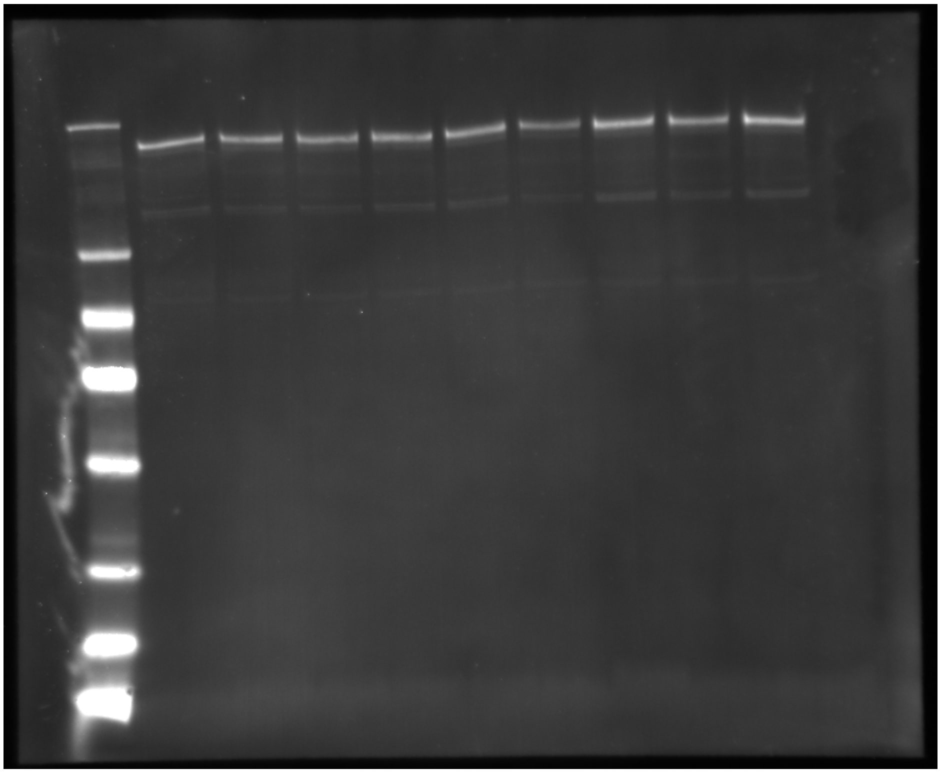


B

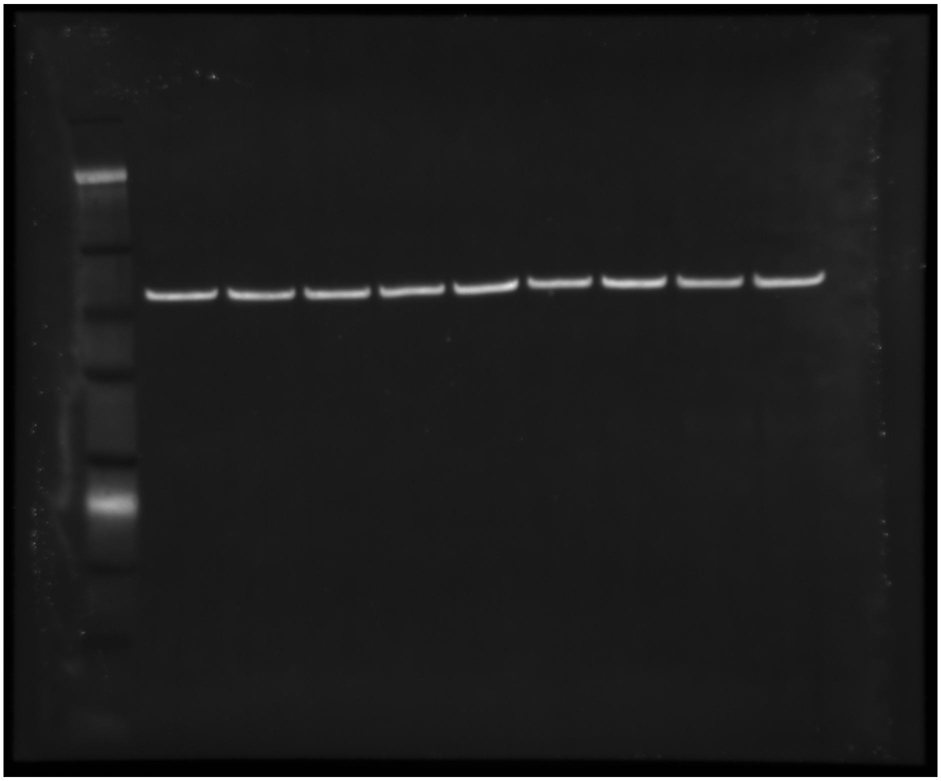


**Supplemental Figure 5.**

A

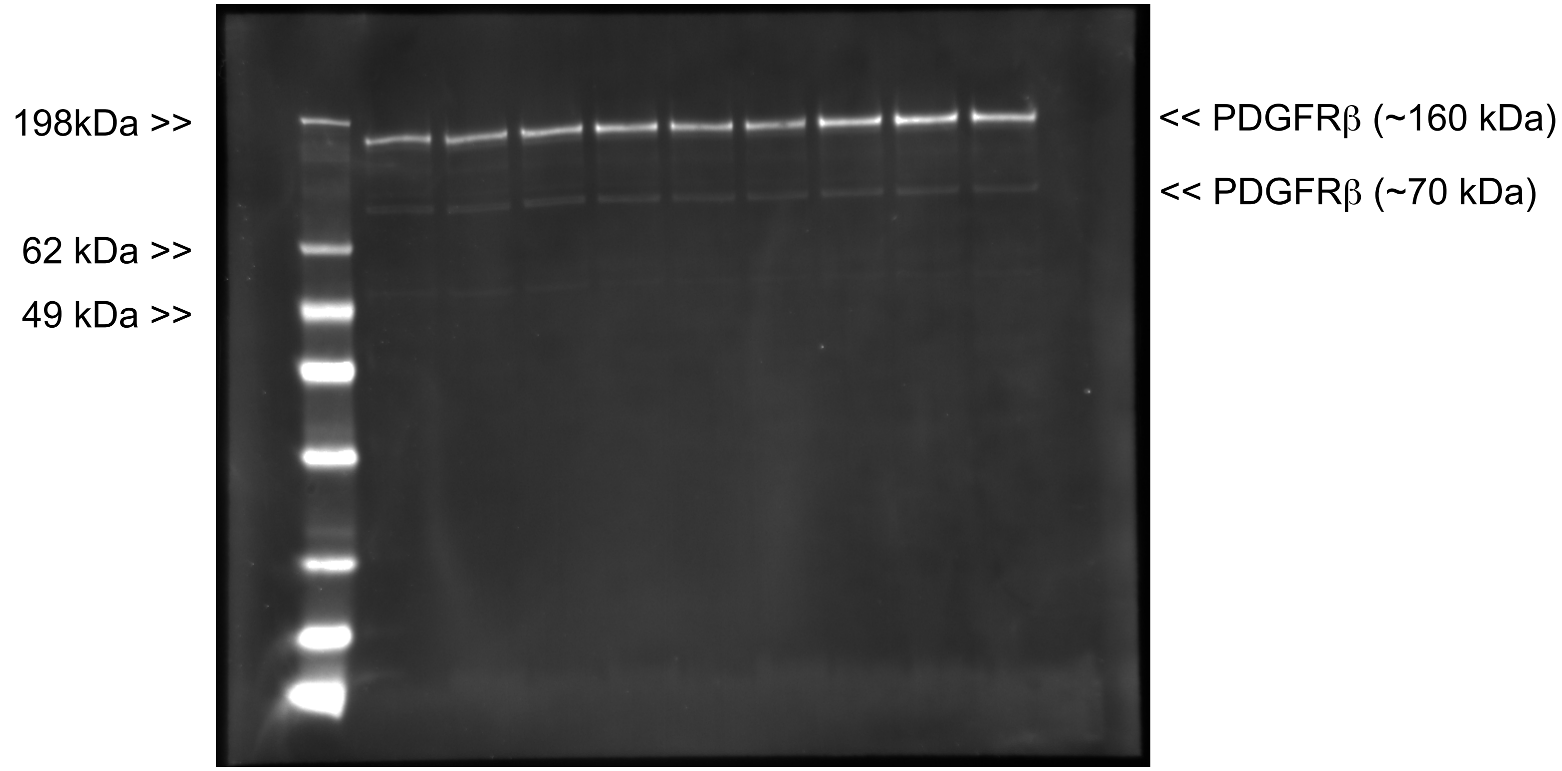


B

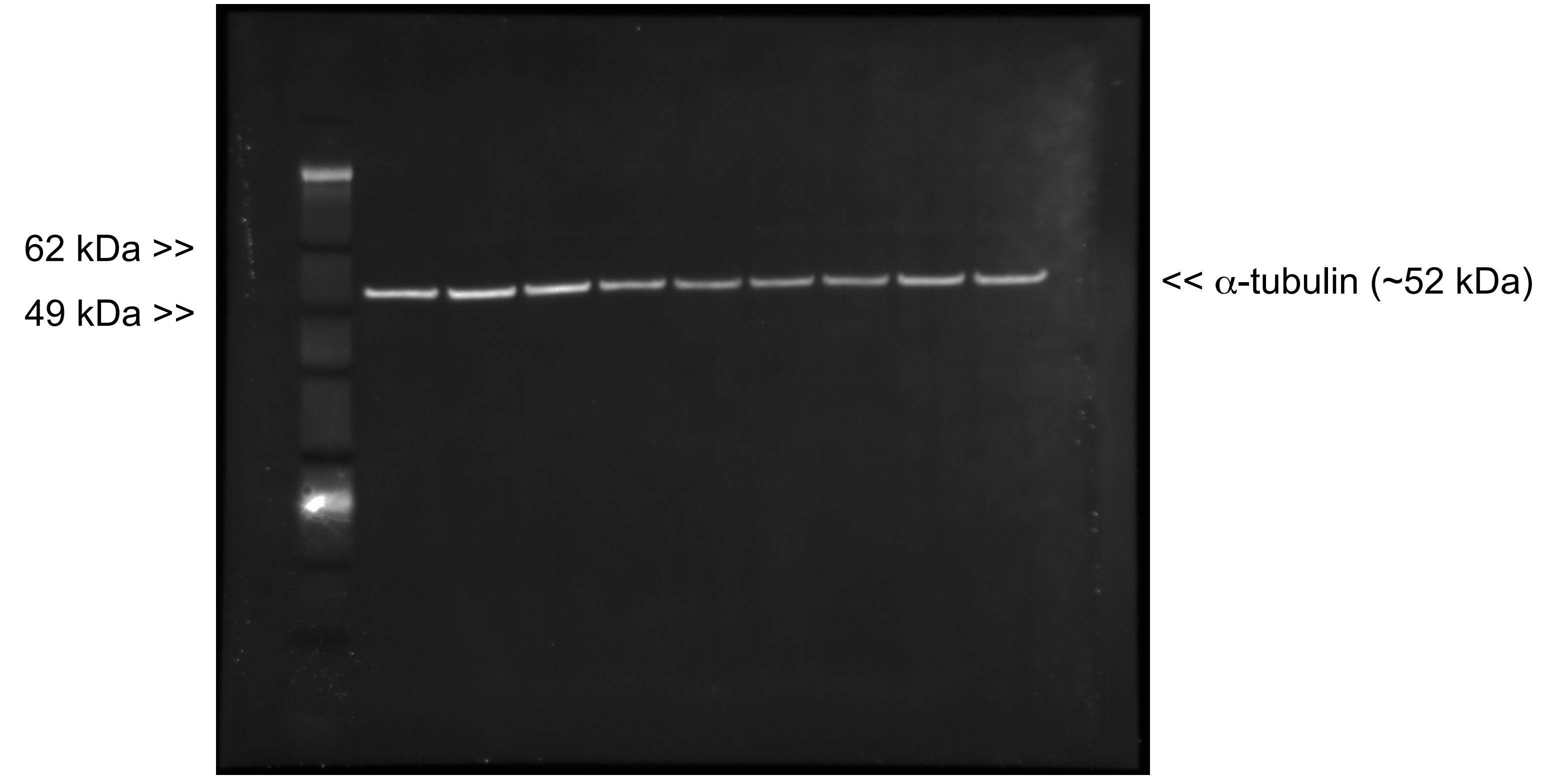


**Supplemental Figure 6.**

**A**

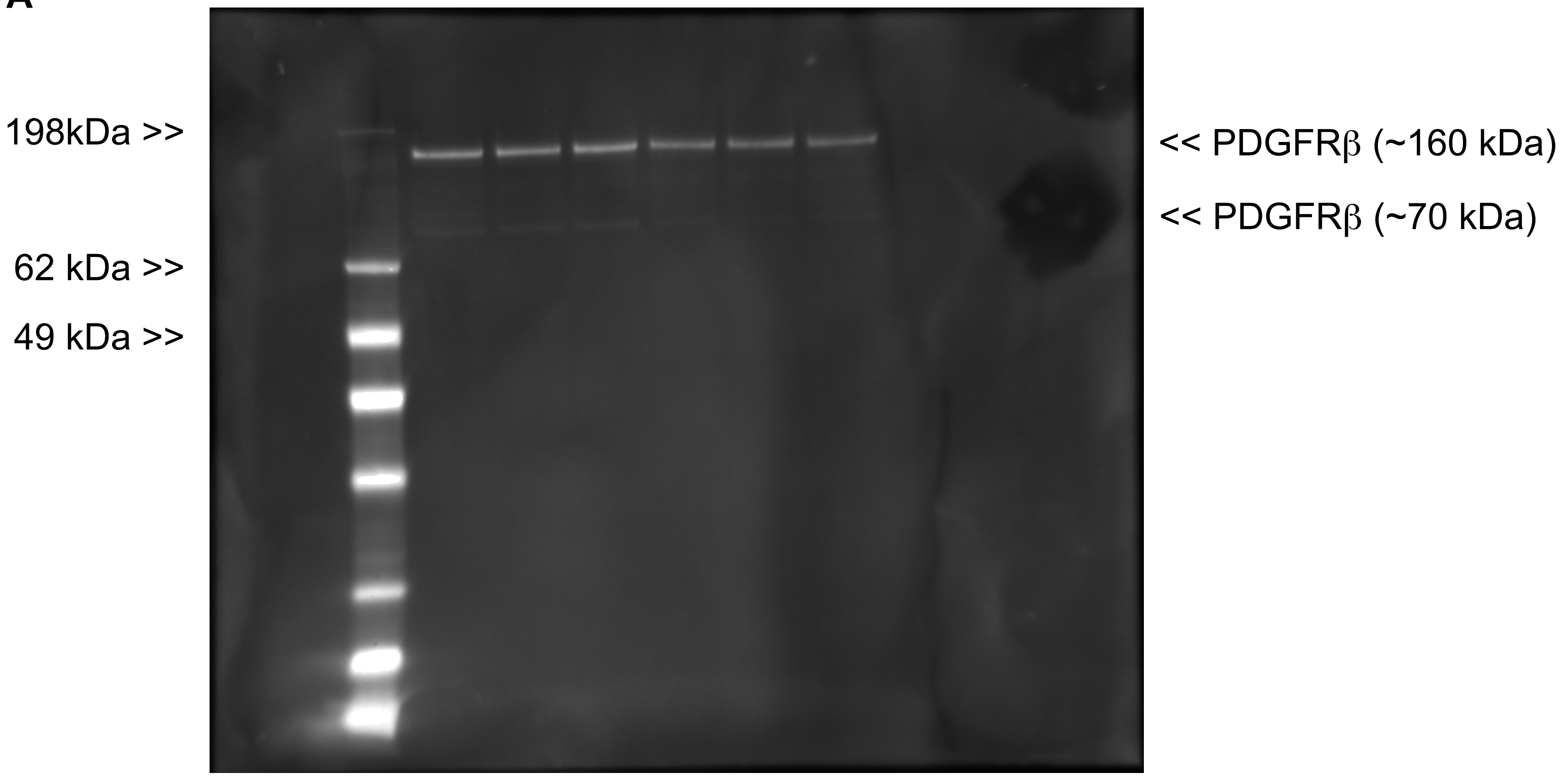


**B**

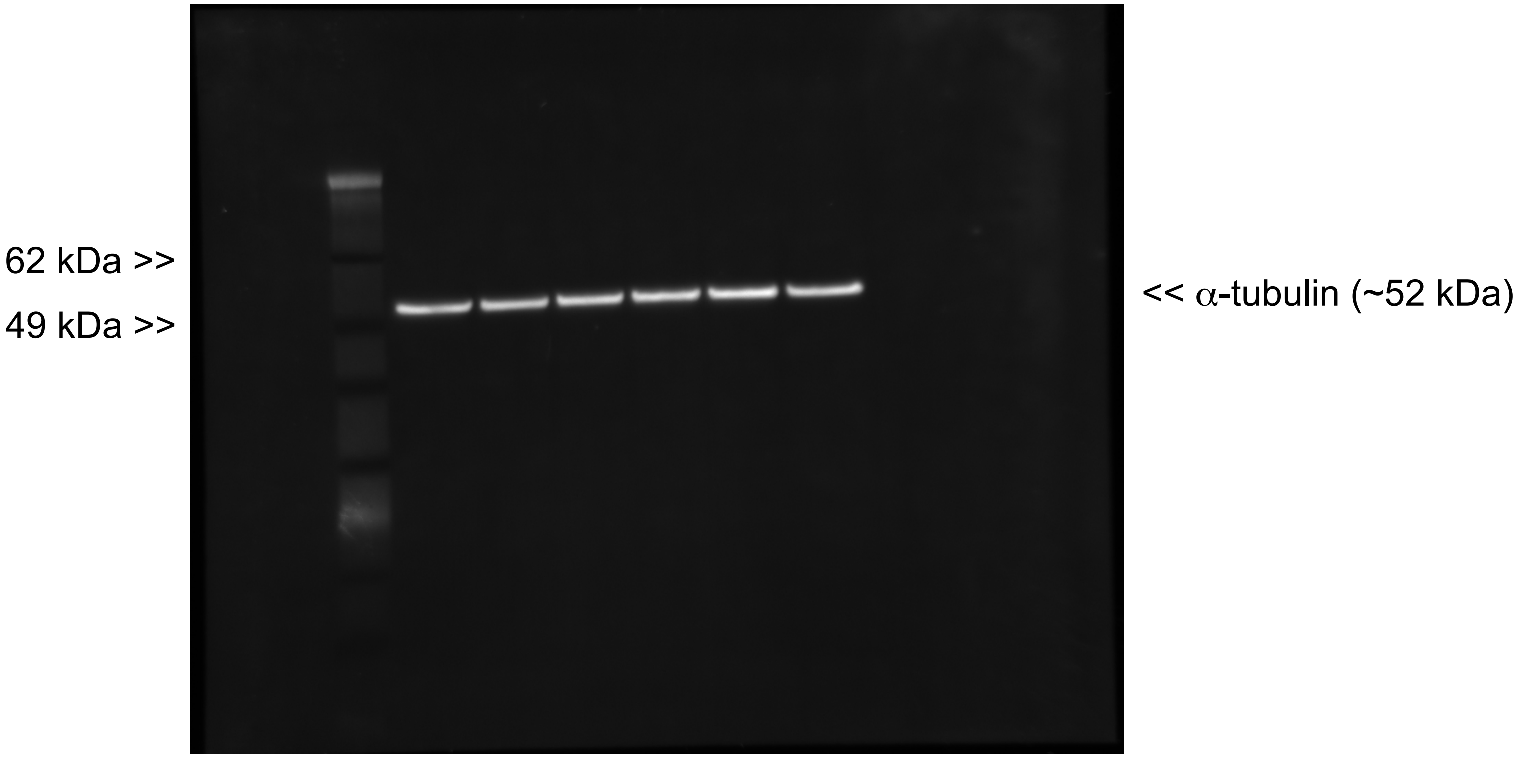


**Supplemental Figure 7.**

**A**

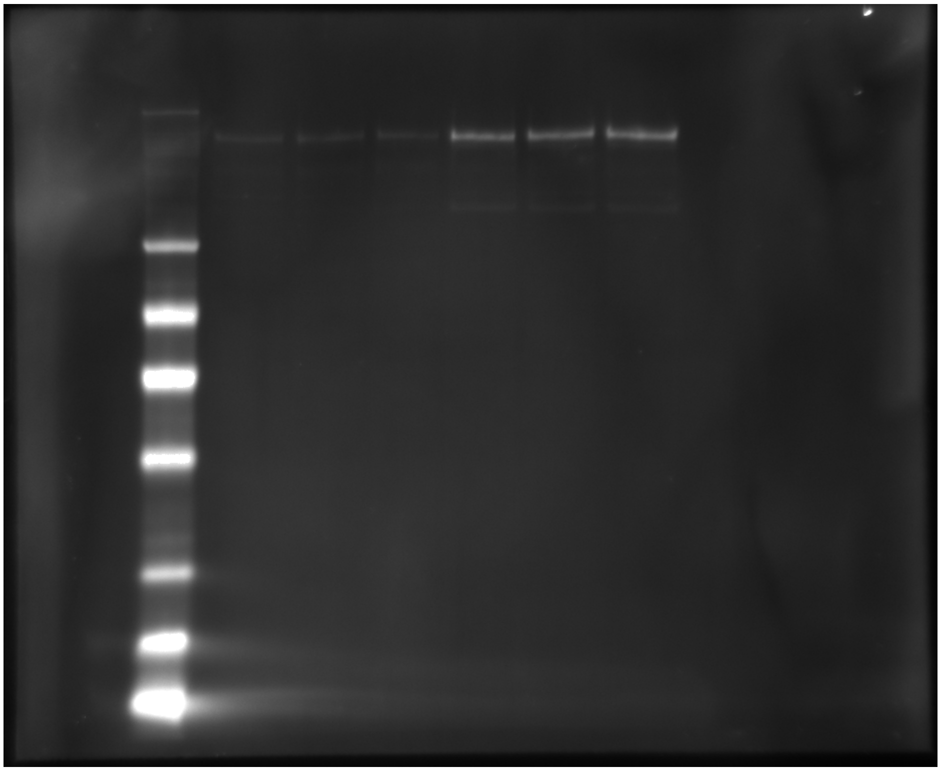


**B**

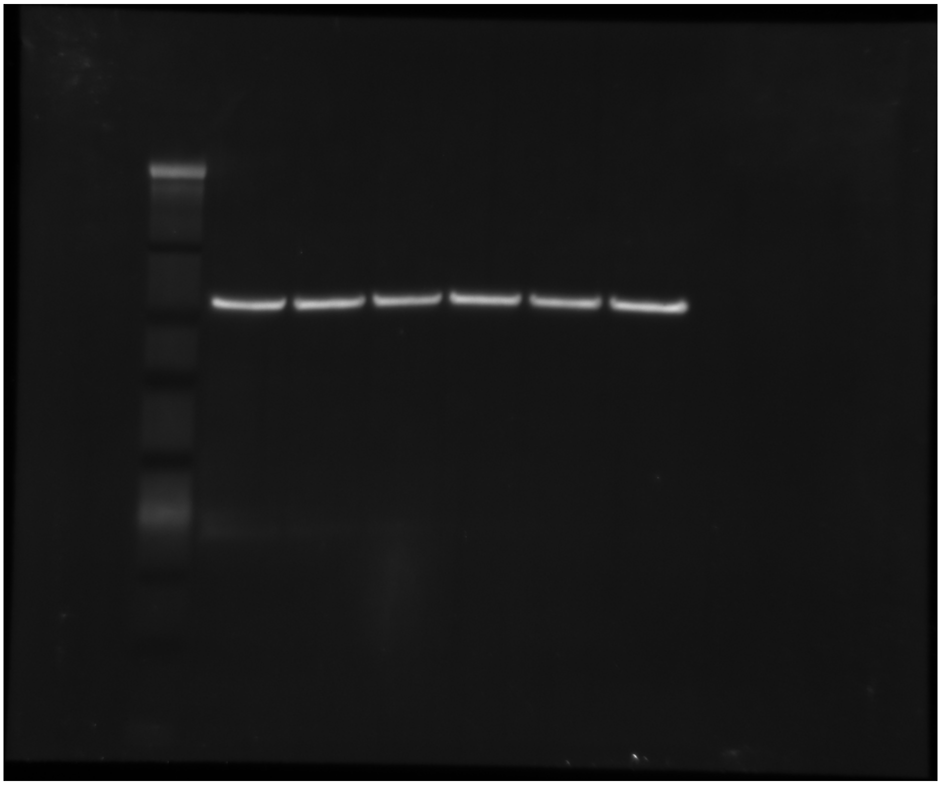


**Supplemental Figure 8.**

**A**

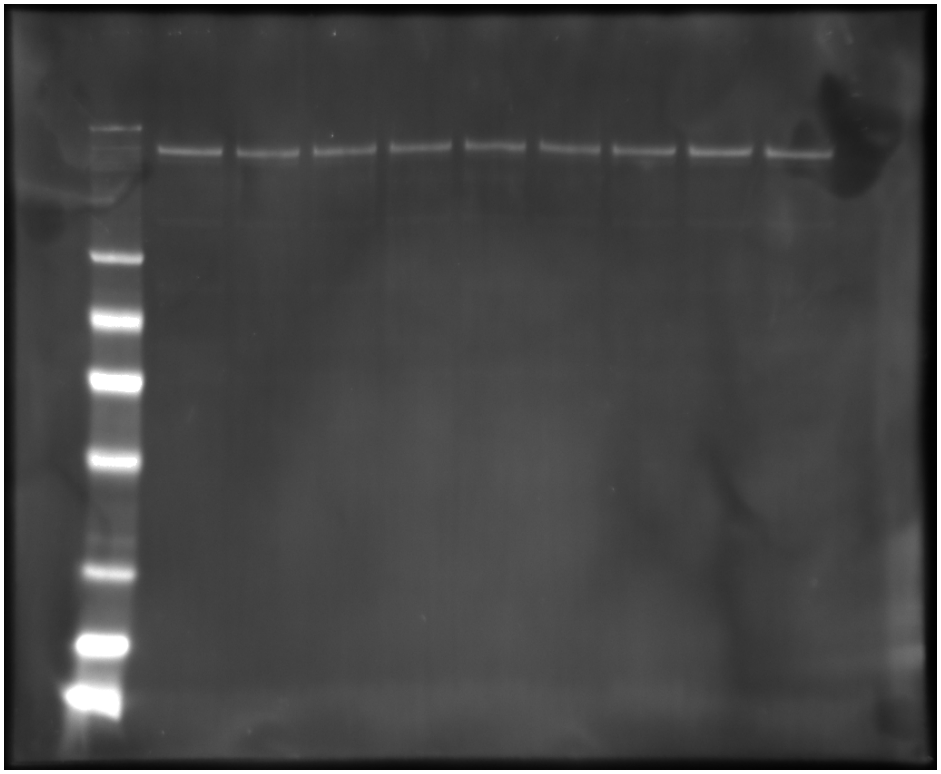


**B**

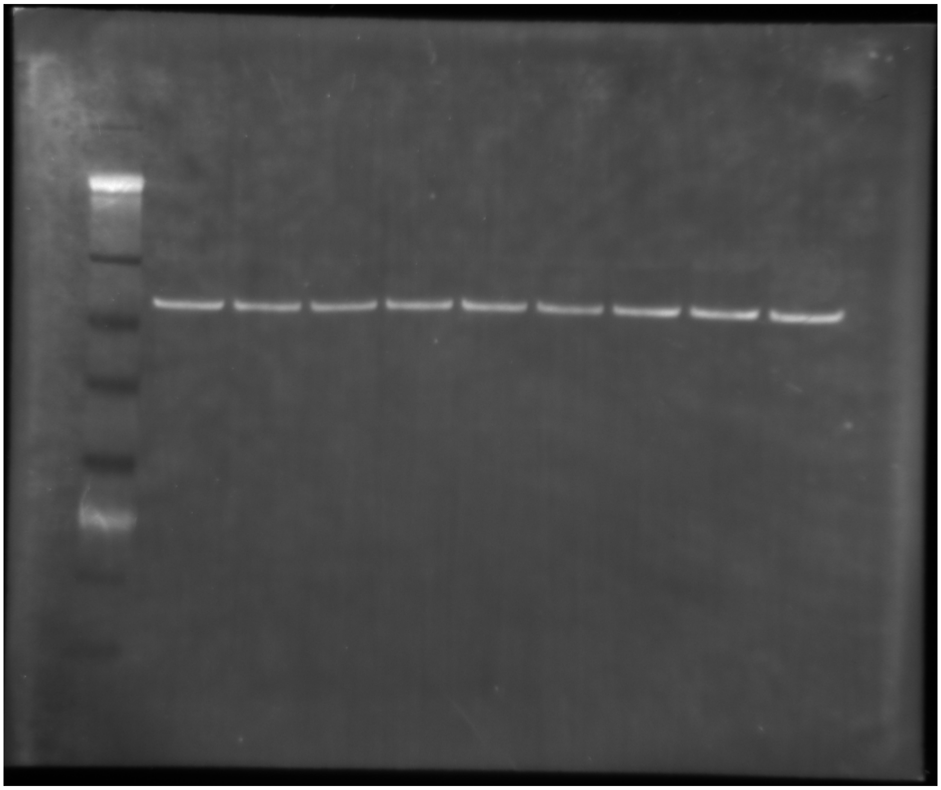


**Supplemental Figure 9.**

A

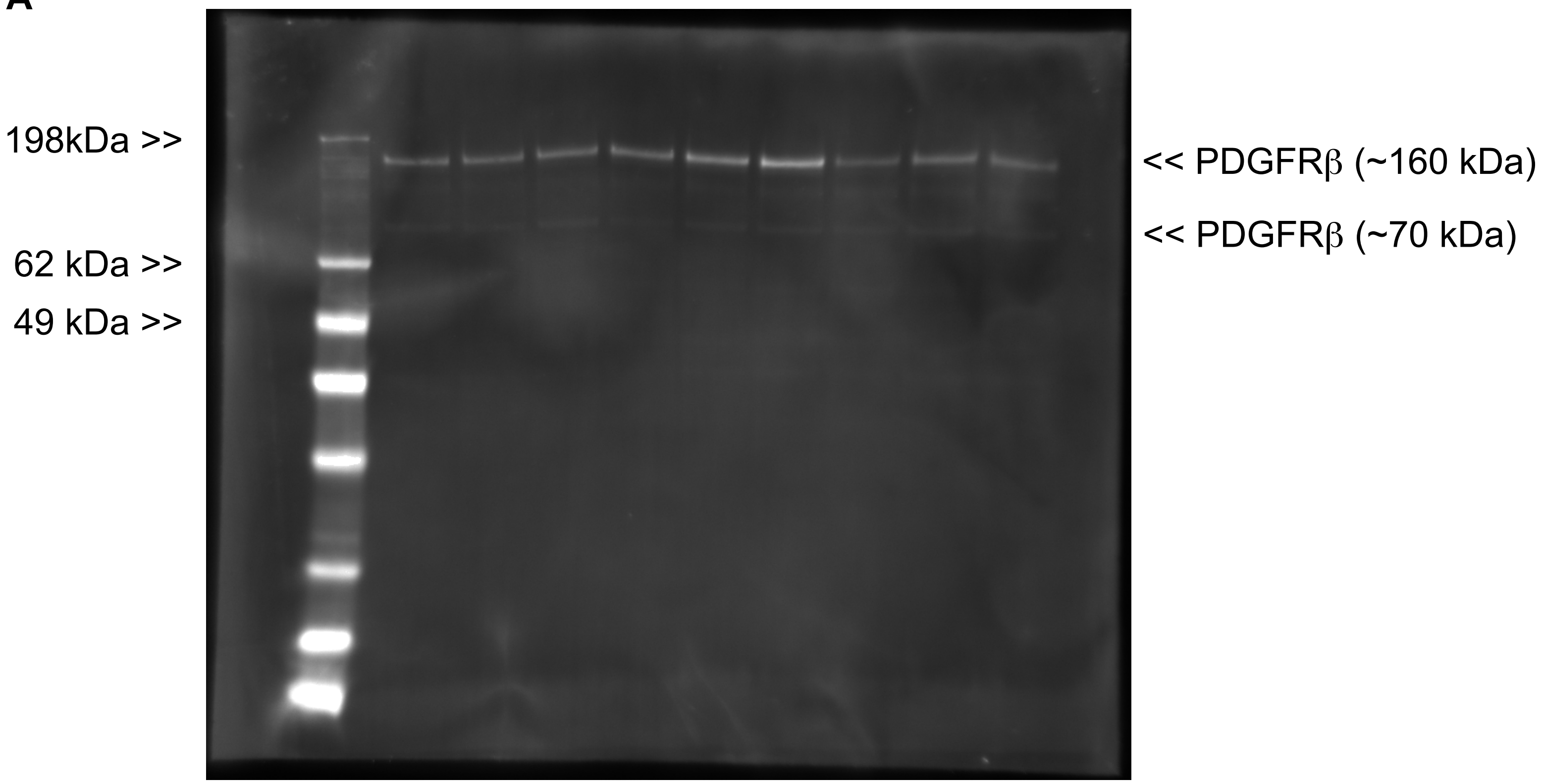


B

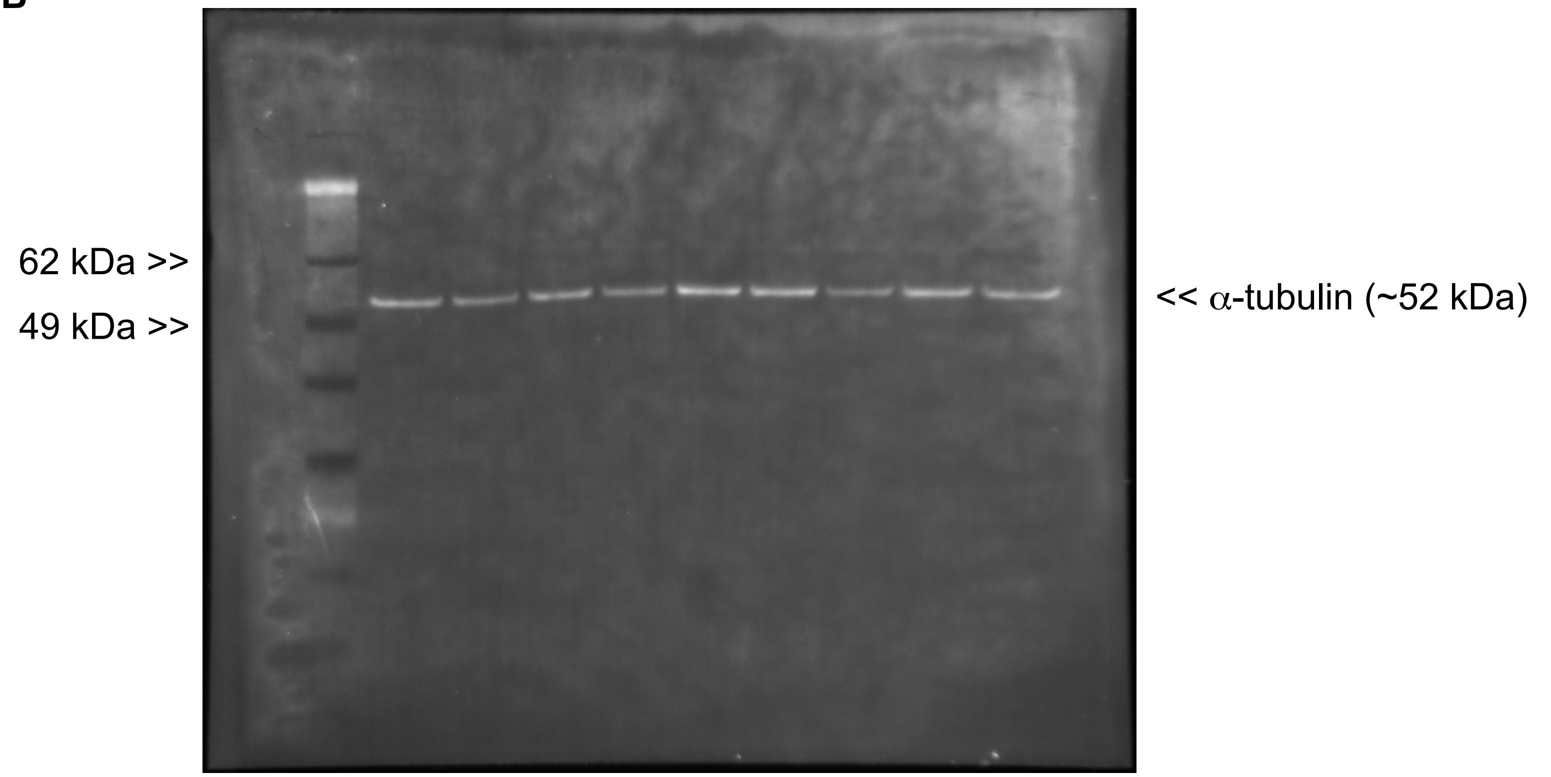


**Supplemental Figure 10.**

A

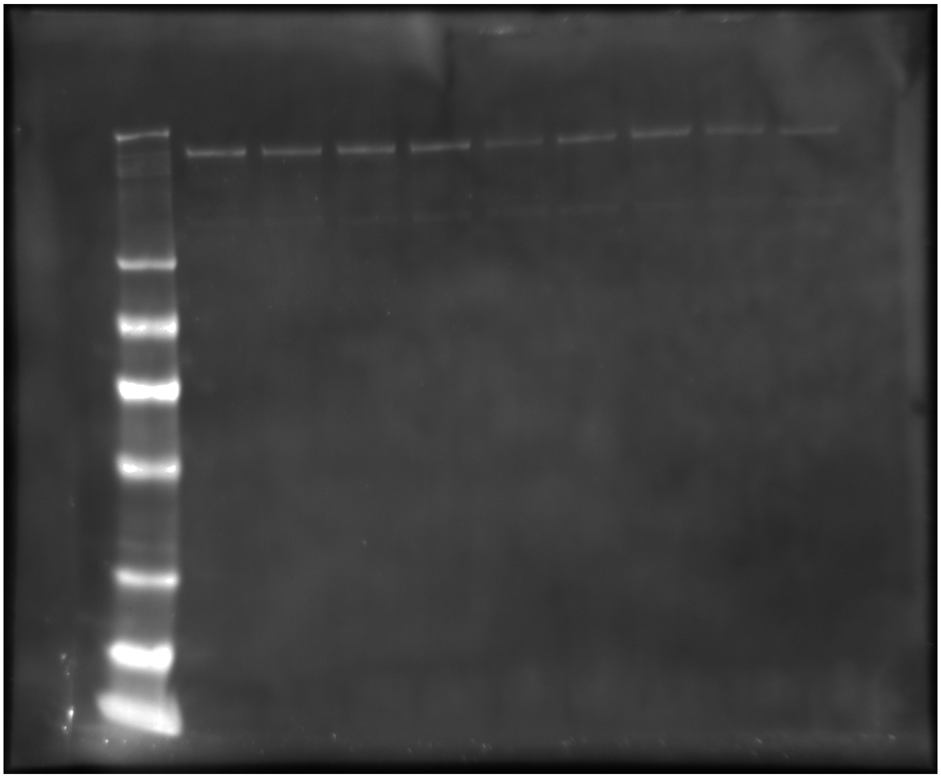


B

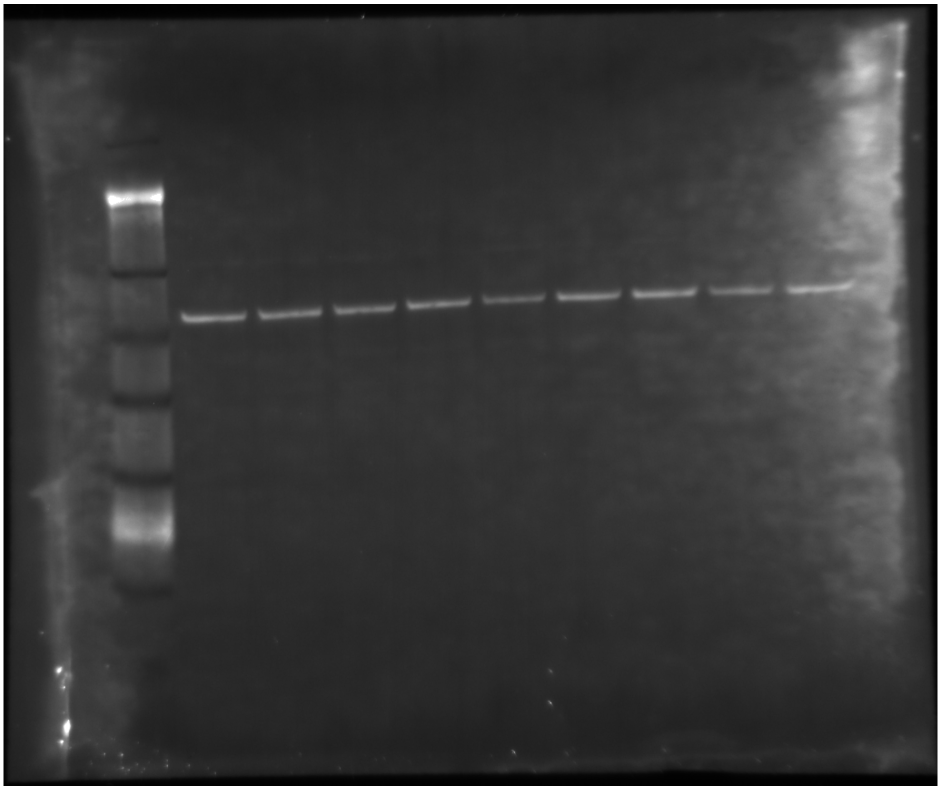


**Supplemental Figure 11.**

A

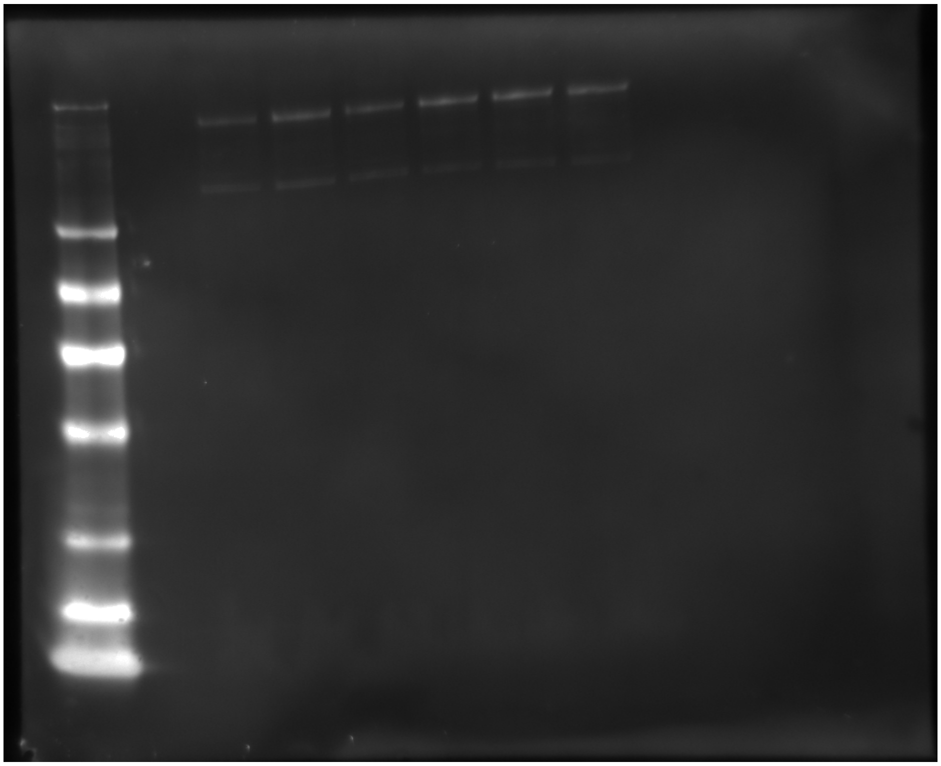


B

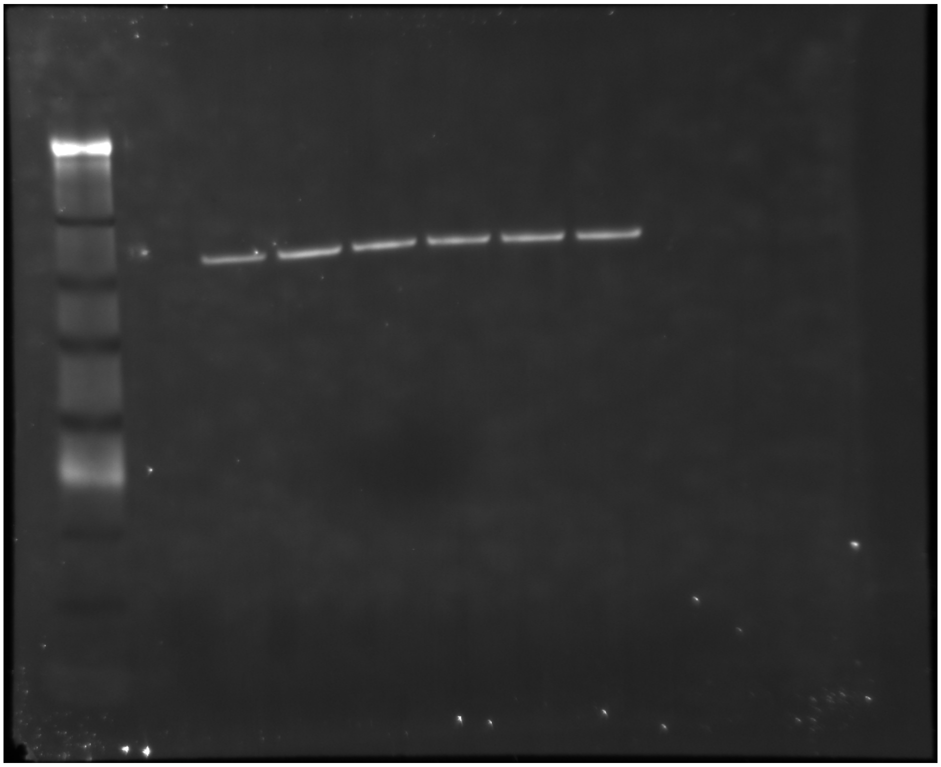


**Supplemental Figure 12.**

A

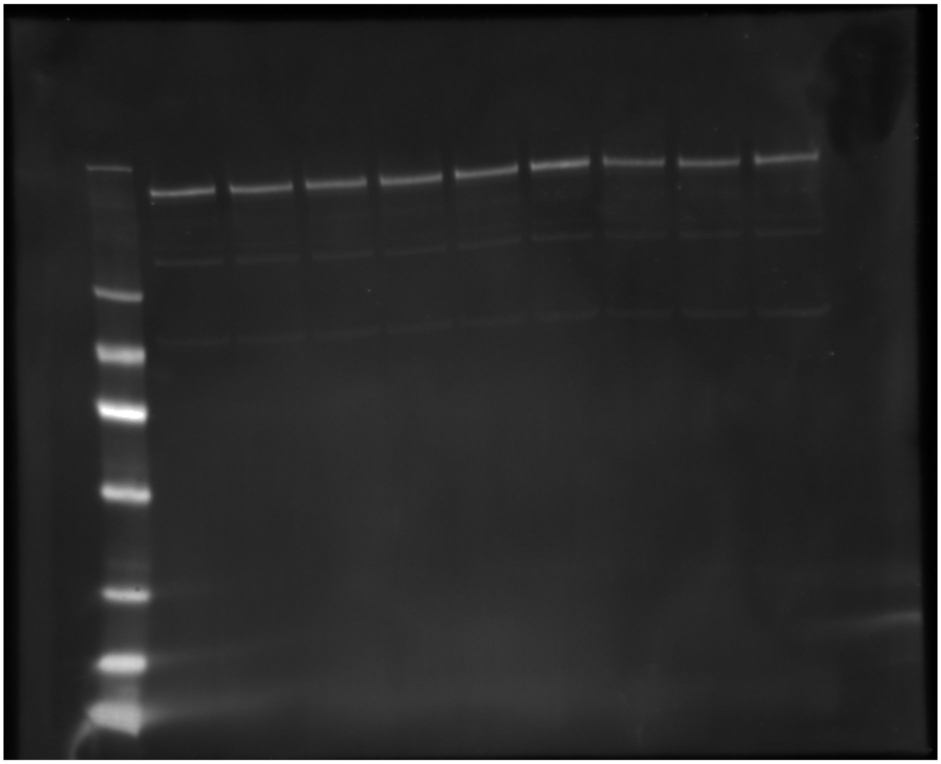


B

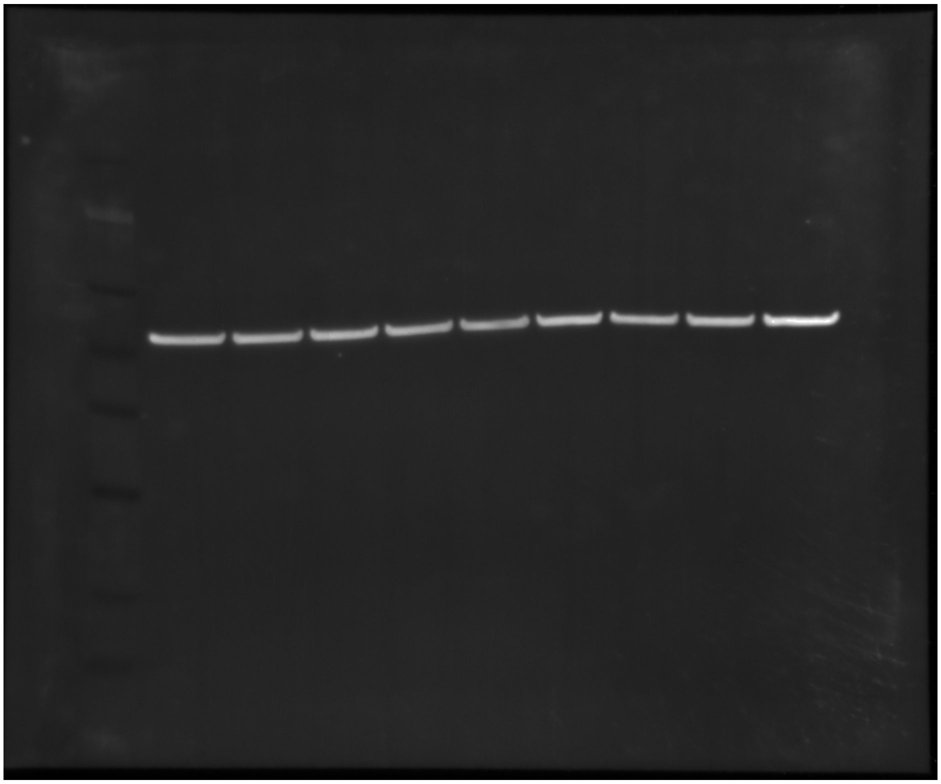


**Supplemental Figure 13.**

**A**

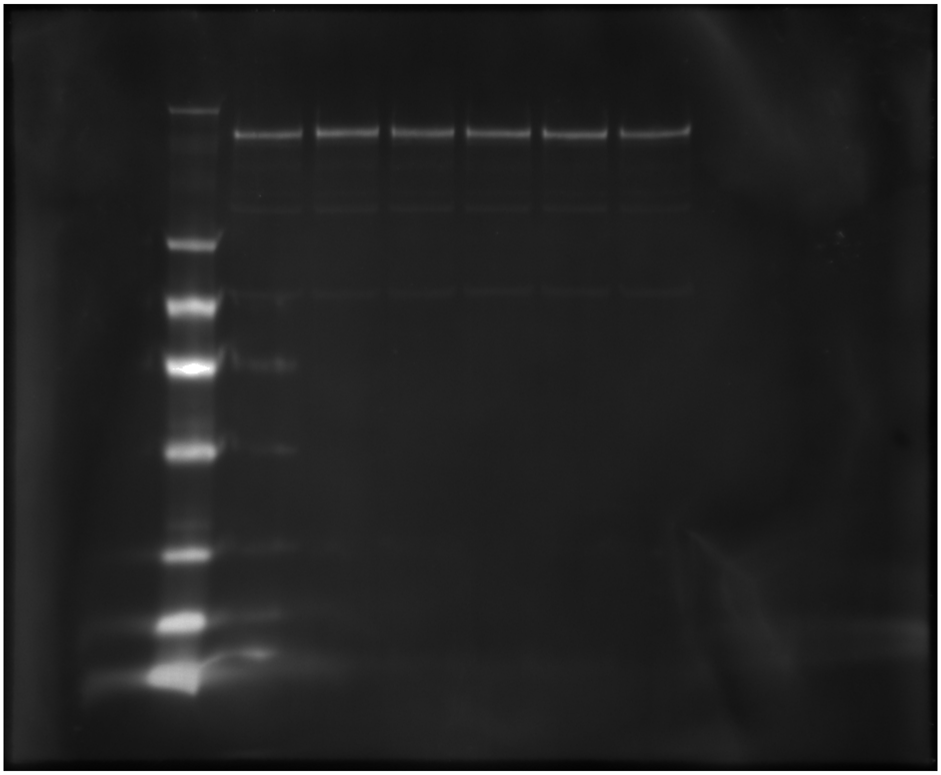


**B**

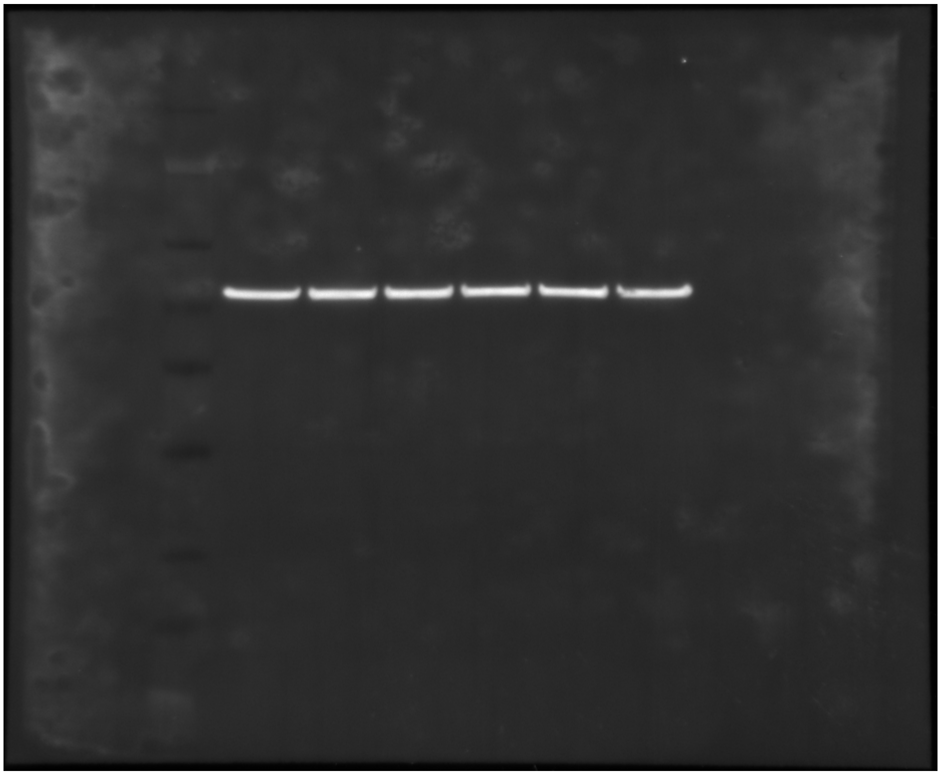


**Supplemental Figure 14.**
